# γ-TuRCs are required for asymmetric microtubule nucleation from the somatic Golgi of *Drosophila* neurons

**DOI:** 10.1101/2021.09.24.461707

**Authors:** Amrita Mukherjee, Paul T. Conduit

## Abstract

Microtubules are polarised polymers nucleated by multi-protein γ-tubulin ring complexes (γ-TuRCs). Within neurons, microtubule polarity is plus-end-out in axons and mixed or minus-end-out in dendrites. Previously we showed that within the soma of *Drosophila* sensory neurons γ-tubulin localises asymmetrically to Golgi stacks, Golgi-derived microtubules grow asymmetrically towards the axon, and growing microtubule plus-ends are guided towards the axon and restricted from entering dendrite in a Kinesin-2-dependent manner (Mukerjee et al., 2020). Here we show that depleting γ-TuRCs perturbs the direction of microtubule growth from the Golgi stacks, consistent with a model for asymmetric microtubule nucleation involving γ-TuRCs and other nucleation-promoting factors. We also directly observe microtubule turning along microtubule bundles and show that depleting APC, proposed to link Kinesin-2 to plus ends, reduces microtubule turning and increases plus end growth into dendrites. These results support a model of asymmetric nucleation and guidance within the neuronal soma that helps establish and maintain overall microtubule polarity.

## Introduction

Microtubules are a key component of the cytoskeleton and play an essential role in neuronal development and function (Stiess and Bradke, 2011). They provide pushing forces for neurite growth and ‘tracks’ for the transport of various cargos between the soma and neurite terminals. Polarity of a microtubule is defined by the α/β-tubulin dimers that join head-to-tail and associate side-by-side during microtubule assembly. α-tubulin is exposed at the microtubule’s minus end and β-tubulin is exposed at the plus end. Plus and minus ends associate with different proteins and display different growth and shrinkage dynamics. Moreover, specific molecular motors (Kinesins and Dynein) ‘read’ the inherent microtubule polarity and ‘walk’ towards either the plus or minus end of the microtubule. In neurons, microtubule polarity differs between axons and dendrites, with axons containing predominantly plus-end-out microtubules and dendrites having either mixed polarity in mammals or predominantly minus-end-out polarity in *Drosophila* and *C. elegans* (Kapitein and Hoogenraad, 2015; Rolls and Jegla, 2015). Correct microtubule polarity within neurons is extremely important as the difference in polarity between axons and dendrites allows the selective transport of molecules into the different compartments, facilitating correct neuronal development and maintenance (Nirschl et al., 2017; Tas et al., 2017). Nevertheless, our understanding of how microtubule polarity within neurons is established and maintained remains incomplete.

Several mechanisms have been shown or proposed to help establish and maintain correct microtubule polarity within neurons, including motor-driven sorting of microtubules (Baas and Lin, 2011; Castillo et al., 2019; Lin et al., 2012; Lu and Gelfand, 2017; Rao et al., 2017; Yan et al., 2013), guidance of growing microtubule plus ends (Mattie et al., 2010; Mukherjee et al., 2020; Weiner et al., 2016), and polarity-dependent differences in microtubule stability (Yau et al., 2016). Another important mechanism that contributes to microtubule polarity within neurons is the regulation of microtubule nucleation (Lüders, 2020). The kinetically unfavourable process of microtubule nucleation can be catalysed by multi-protein γ-tubulin ring complexes (γ-TuRCs) (Farache et al., 2018; Kollman et al., 2011; Tovey and Conduit, 2018). The process of γ-TuRC-mediated nucleation is inherently polar as the α-tubulin molecules from α/β-tubulin dimers associate with the γ-tubulin molecules of a γ-TuRC. After formation and stabilisation of an initial seed, microtubule polymerisation continues at the microtubule plus end, which grows away from the γ-TuRC. Thus, the γ-TuRC remains at the microtubule minus end while the plus end grows outwards, providing a way to regulate microtubule polarity when γ-TuRCs are anchored to specific intracellular sites.

Despite a clear role for γ-TuRCs in microtubule nucleation, other microtubule associated proteins, particularly TOG-domain and CAMSAP proteins, can be important for efficient microtubule nucleation and can also promote microtubule nucleation independently of γ-TuRCs both *in vitro* and *in vivo* (Coquand et al., 2021; Flor-Parra et al., 2018; Hannak et al., 2002; Hussmann et al., 2016; King et al., 2020; Llamazares et al., 1999; Nashchekin et al., 2016; Popov et al., 2002; Roostalu et al., 2015; Sampaio et al., 2001; Slep and Vale, 2007; Tanaka et al., 2012; Thawani et al., 2018; Tsuchiya and Goshima, 2021; Wang et al., 2015; Wieczorek et al., 2015; Woodruff et al., 2017; Zheng et al., 2020). These proteins bind directly to α/β-tubulin dimers and can either catalyse their assembly into microtubule polymers or stabilise nascent microtubule polymers. The extent to which γ-TuRC-independent nucleation occurs within cells remains unclear, but examples of γ-TuRC independent nucleation occurring both naturally (Coquand et al., 2021; Nashchekin et al., 2016; Tanaka et al., 2012; Wang et al., 2015; Zheng et al., 2019) and after γ-TuRC depletion (Hannak et al., 2002; Llamazares et al., 1999; Sampaio et al., 2001; Tsuchiya and Goshima, 2021) have been reported.

Microtubule nucleation normally occurs at specific sites called microtubule organising centres (MTOCs) (Paz and Lüders, 2018; Sanchez and Feldman, 2016; Tillery et al., 2018), where microtubule associated proteins accumulate and where γ-TuRCs are thought to be activated. The centrosome functions as a major MTOC in cycling cells (Conduit et al., 2015), but several other MTOCs have been identified in differentiated cells, including neurons (Paz and Lüders, 2018; Sanchez and Feldman, 2016). In cultured mammalian neurons, the centrosome is deactivated during early development (Stiess et al., 2010) and has no apparent MTOC activity or functional necessity within *Drosophila* neurons (Nguyen et al., 2011).

In recent years it has become clear that non-centrosomal MTOCs (nc-MTOCs) play an important role in organising the microtubule cytoskeleton within neurons. Centrosome-like protein accumulations function as MTOCs at the dendritic tips of ciliated sensory neurons in *C. elegans* (Garbrecht et al., 2021; Harterink et al., 2018; Magescas et al., 2021). Fragments of Golgi known as Golgi outposts have been proposed to function as MTOCs within the dendrites of *Drosophila* dendritic arborisation (da) neurons and within mouse oligodendrocytes (Fu et al., 2019; Ori-McKenney et al., 2012; Valenzuela et al., 2020; Yalgin et al., 2015; Yang and Wildonger, 2020; Zhou et al., 2014). While Golgi outposts are important for microtubule polarity in oligodendrocytes (Fu et al., 2019), they are not essential within *Drosophila* dendrites (Nguyen et al., 2014), although mis-positioning these Golgi outposts into axons does perturb axonal microtubule polarity (Arthur et al., 2015; Yang and Wildonger, 2020). Interestingly, γ-TuRCs do not accumulate at Golgi outposts in oligodendrocytes (Fu et al., 2019) or at the vast majority of Golgi outposts in da neurons (Mukherjee et al., 2020), suggesting an alternative molecular pathway for microtubule generation. Indeed, tubulin polymerising promoting protein (TPPP) promotes nucleation from Golgi outposts in mouse oligodendrocytes (Fu et al., 2019). In addition to Golgi outposts, endosomes have recently been identified as MTOCs at dendritic branchpoints of *Drosophila* da neurons (Weiner et al., 2020) and at the dendritic tips of *C. elegans* PVD neurons (Liang et al., 2020). These endosomes do recruit γ-TuRCs and are essential for maintaining microtubule polarity within the dendrites of both species. Microtubule nucleation can also occur from the sides of preexisting microtubules via Augmin/HAUS-mediated recruitment of γ-TuRCs, and this mode of nucleation is important in the axons and dendrites of mammalian hippocampal neurons (Cunha-Ferreira et al., 2018; Qu et al., 2019; Sánchez-Huertas et al., 2016). Augmin/HAUS-mediated nucleation generates new microtubules that have the same polarity as the original ‘mother’ microtubule (Kamasaki et al., 2013; Petry et al., 2013), explaining why it can be important for maintaining microtubule polarity within neurons (Cunha-Ferreira et al., 2018; Qu et al., 2019; Sánchez-Huertas et al., 2016). In the basal processes of mammalian and human radial glial cells, microtubules are generated at neurite expansions (varicosities) that lack γ-tubulin but that are filled with the minus end binding CAMSAP proteins (Coquand et al., 2021). Similar varicosities have been observed in class I *Drosophila* da neurons (Mukherjee et al., 2020), but these are filled with γ-tubulin and other centrosomal proteins and it remains to be determined if they organise microtubules. Disrupting Augmin/HAUS-mediated microtubule nucleation in hippocampal neurons and CAMSAP-mediated microtubule organisation in radial glial cells disrupts microtubule polarity (Coquand et al., 2021; Cunha-Ferreira et al., 2018; Qu et al., 2019; Sánchez-Huertas et al., 2016). In summary, various modes of microtubule nucleation have been discovered both across species and within the same neuron of the same species and play a role, to differing degrees, in regulating microtubule polarity within neurons.

Along with regulating microtubule nucleation, guiding the direction of microtubule plus end growth also contributes to microtubule polarity control. Within *Drosophila* class I da neurons, plus end turning at dendritic branch points can potentially occur towards or away from the soma, but the vast majority turn towards the soma, adding to the predominantly minus-end-out microtubule polarity within these dendrites (Mattie et al., 2010). This is dependent on the plus-end-directed motor Kinesin-2 (Mattie et al., 2010). Kinesin-2 motors have diverse functions and can exist in different configurations, including a heterotrimeric complex of two motor domains and one regulatory kinesin associated protein 3 (KAP3) (Scholey, 2012). In addition to transporting cargo along microtubules, Kinesin-2 has been proposed to associate with the plus ends of microtubules and guide their growth along other neighbouring microtubules (Mattie et al., 2010). The regulatory component of Kinesin-2, KAP3, interacts with Adenomatous polyposis coli (APC) (Jimbo et al., 2002; Mattie et al., 2010), which can accumulate at the plus ends of microtubules (Mimori-Kiyosue et al., 2000; Näthke et al., 1996) and interact with End Binding Protein 1 (EB1) (Mattie et al., 2010). Thus, a reasonable model is that APC can link Kinesin-2 to the growing plus ends of microtubules via an interaction with EB1 in such a manner that the motor domains of Kinesin-2 remain free to exert a ‘walking’ force along adjacent microtubules and direct the growing plus end towards the plus end of the adjacent microtubules (Mattie et al., 2010). This model is supported by *in vitro* experiments showing that artificially tethering Kinesin-2 to the plus ends of growing microtubules is sufficient to induce microtubule turning along adjacent microtubules (Chen et al., 2014; Doodhi et al., 2014).

We recently examined the behaviour of microtubules within the soma of *Drosophila* class I da neurons and proposed that microtubule polarity within axons and dendrites could be regulated by controlled microtubule nucleation and plus end guidance within the soma (Mukherjee et al., 2020). There are four classes of da neurons ranging from proprioceptive class I neurons, with the simplest dendritic arbor morphology, to the nociceptive class IV neurons, which have large highly branched dendritic arbors (Grueber et al., 2002). We showed that γ-tubulin accumulates asymmetrically at the cis-face of the multiple Golgi stacks that are distributed throughout the soma of both class I and class IV da neurons and the soma of other neighbouring sensory neurons, including the external sensory (ES) neurons (Mukherjee et al., 2020). Using class I da neurons as a model and EB1-GFP as a marker of growing microtubules we found that many microtubule growth events originated from the somatic Golgi stacks, including after cold-induced microtubule depolymerisation. Intriguingly, the initial direction of growth of these microtubules was preferentially towards the axon, suggestive of asymmetric microtubule nucleation. The vast majority of the plus ends of these microtubules also turned towards the axon entry site and away from dendritic entry sites within the soma. These turning events were dependent on Kinesin-2, similar to the turning events observed previously within branchpoints (Mattie et al., 2010; Weiner et al., 2016). Based on our observations, we proposed that plus-end-associated Kinesin-2 guides growing microtubule plus ends along a polarised microtubule network within the soma that is generated, at least in part, by asymmetric nucleation from the somatic Golgi (Mukherjee et al., 2020). In addition to Kinesin-2-mediated turning, we also showed that the majority of microtubule plus ends that did grow towards a dendrite failed to enter, which was also dependent on Kinesin-2. We proposed that plus-end-associated Kinesin-2 restricts entry into dendrites because it’s motor domains encounter and attempt to walk back along dendritic microtubules that have the opposite polarity to the growing microtubule. Nevertheless, we had yet to establish a role for γ-tubulin in microtubule nucleation at the somatic Golgi stacks or to show directly that microtubules turn along other microtubules within the soma.

In this paper, we show that depletion of the essential γ-TuRC component Grip91^GCP3^ from da neurons results in the loss of endogenously-regulated γ-tubulin-GFP from the somatic Golgi stacks, showing that Golgi-associated γ-tubulin-GFP is part of multiprotein γ-TuRCs. By using EB1-GFP comets to track microtubule growth events we find that microtubules can still originate from the Golgi stacks after γ-TuRC depletion, suggesting that nucleation can be promoted by other microtubule associated proteins. Nevertheless, we find that these Golgi-derived microtubules no longer preferentially grow towards the axon, suggesting that the primary role of γ-TuRCs at somatic Golgi stacks is to orient rather than initiate microtubule nucleation. In addition, by examining EB1-GFP comets in the presence of a microtubule shaft marker we directly observe microtubules turning along pre-existing microtubule bundles within the soma. We show that turning along microtubule bundles close to the cell periphery occurs preferentially towards the axon, supporting the idea that a polarised microtubule network exists within the soma and helps guide growing microtubules towards the axon. We also show that depletion of APC reduces the proportion of growing microtubules that turn within the soma and increases the proportion that enter dendrites, supporting our model in which APC links Kinesin-2 to microtubule plus ends and thus facilitates plus end guidance within the neuronal cell body.

## Results

### RNAi depletion of Grip91^GCP3^ strongly perturbs dendritic branching of pupal class IV da neurons

We began by trying to identify a UAS-driven RNAi construct that would most efficiently deplete γ-TuRCs from da neurons. To establish an assay to examine the efficiency of γ-TuRC RNAi in da neurons we decided to analyse the dendritic branching pattern of class IV v’ada neurons within developing pupae. Dendritic branching of class IV da neurons is particularly sensitive to perturbation of microtubule-related processes (Chen et al., 2012; Norkett et al., 2020; Ori-McKenney et al., 2012; Sears and Broihier, 2016; Wang et al., 2019) and quantifying dendritic arbor morphology under RNAi conditions requires simple genetic and imaging conditions. Class IV da neurons undergo dendritic remodelling at the larval to pupal transition when the dendrites are completely degraded (Williams and Truman, 2005). These neurons then entirely regrow their dendritic arbor during pupal development (Satoh et al., 2012; Shimono et al., 2009). In our hands, UAS-controlled RNAi lines have a stronger effect on dendritic morphology at pupal stages compared to larval stages (unpublished observations, see also below), presumably because in pupae the RNAi construct is effective from the very beginning of dendritic arbor formation. Thus, analysing dendritic arbor morphology in pupae rather than larvae provides a clearer phenotypic readout.

Each γ-TuRC contains 14 γ-tubulin molecules held in a single-turn helical conformation by so-called γ-tubulin complex proteins (GCPs) (Zupa et al., 2021). Of the known γ-TuRC proteins, only γ-tubulin, GCP2 and GCP3 are conserved in all Eukaryotes and are essential for viability, including in *Drosophila* (Tovey and Conduit, 2018). We therefore tested RNAi lines targeting either γ-tubulin, GCP2 or GCP3. Multiple UAS-controlled RNAi constructs are available for each gene but the depletion efficiency of RNAi is unpredictable. We therefore chose to test two independent RNAi lines for each of the three essential γ-TuRC genes (Table S1). There are two γ-tubulin genes in *Drosophila*, a maternally expressed form called γ-tubulin37C and a zygotically expressed form called γ-tubulin23C. RNAi targeting γ-tubulin37C can therefore be used as a negative control within da neurons (Feng et al., 2019; Mukherjee et al., 2020). The names of GCP proteins vary between species making nomenclature confusing. Therefore, throughout this manuscript, we use the *Drosophila* names with the human GCP nomenclature in superscript i.e. Grip84^GCP2^ and Grip91^GCP3^ for *Drosophila* GCP2 and GCP3, respectively.

We used ppk-Gal4 to express the RNAi constructs and measured both the coverage area of the dendritic arbor and the number of dendrite tips at 72h after pupae formation (APF), a time at which the dendritic arbor has grown to a reasonable size for imaging and analysis (Figure 1A). As expected, we found variation in the degree to which arbor area and tip number were affected when using the different RNAi constructs (Figure 1B-D). While each of the two RNAi constructs against γ-tubulin23C and Grip84^GCP2^ resulted in a reduction for both arbor area and dendrite tip number, we found that the strongest reduction occurred when depleting γ-TuRCs using the Grip91^GCP3^-RNAi1 line corresponding to VDRC KK library construct 104667 (hereafter Grip91^GCP3^-RNAi) (Figure 1B-D). We verified by PCR that this line did not contain an aberrant insertion at genomic position 40D (data not shown), which is present in some KK library constructs and can result in unspecific phenotypes (see information on VDRC website). We therefore chose to use this Grip91^GCP3^-RNAi line in the experiments that follow.

**Figure 1.**
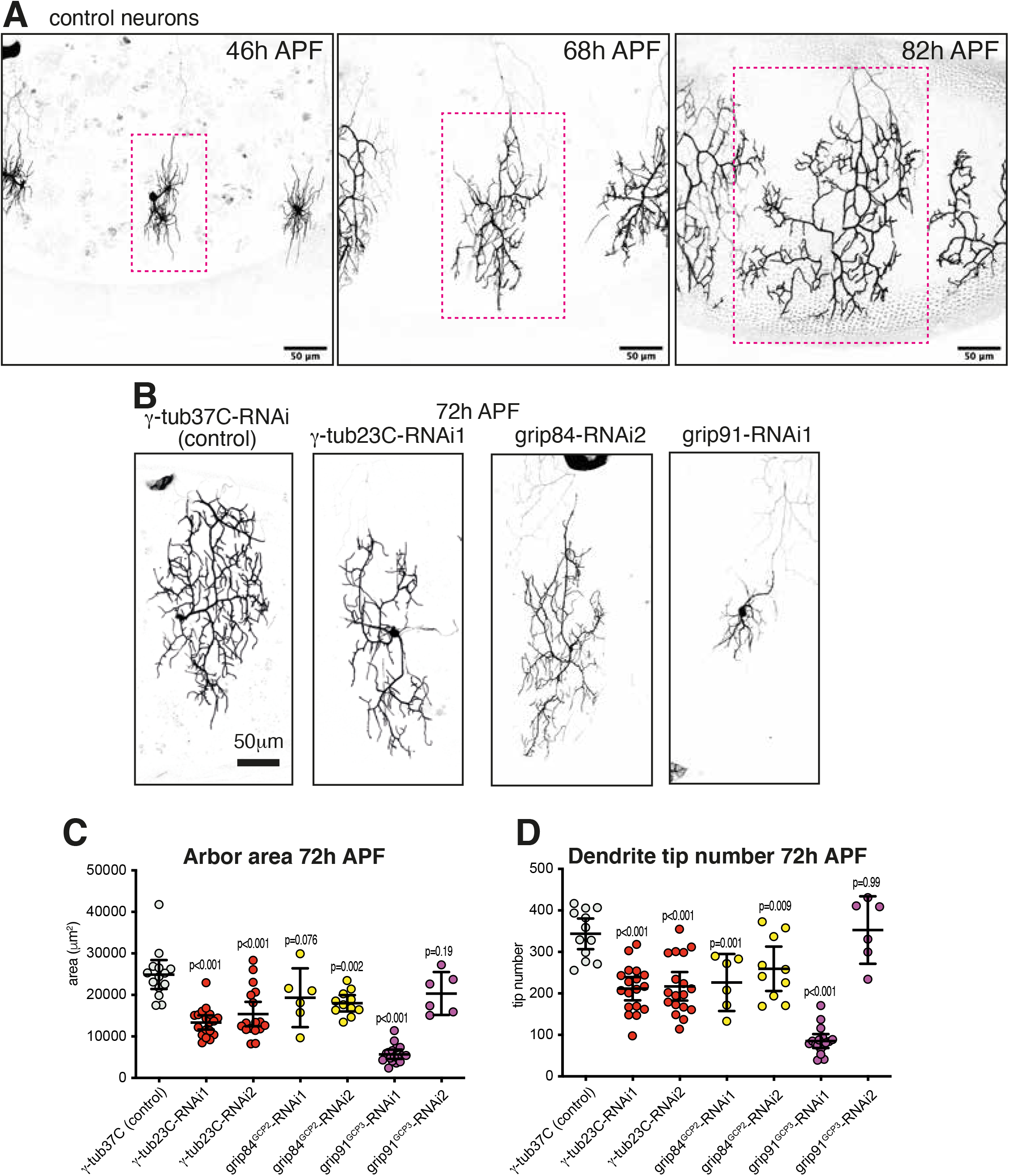
Effect of different RNAi lines targeting essential γ-TuRC components on the dendritic morphology of pupal class IV v’ada neurons. (**A**) Confocal images show class IV v’ada neurons in control flies expressing CD4-tdGFP at different times after pupal formation (APF), as indicated. Red dotted line indicates a single dendritic field. Note that the neurons are from different animals but imaged within the same abdominal segment. (**B**) Confocal images show class IV v’ada neurons within flies expressing CD4-tdGFP, UAS-Dicer2 and different UAS-RNAi lines, as indicated, at 72h after pupal formation (APF). (**C,D**) Graphs show the dendritic arbor area (μm^2^) (C) or the number of dendritic tips (D) of class IV v’ada neurons within flies expressing CD4-tdGFP, UAS-Dicer2 and different UAS-RNAi lines, as indicated, at 72h after pupal formation (APF). Each point on the graph represents a different neuron. Mean and 95% CI’s are indicated. Scale bars are indicated within the images. Note how expressing most of the RNAi constructs reduces arbor area and tip number but that expressing Grip91^GCP3^-RNAi1 results in the strongest reduction in both indicators. Grip91^GCP3^-RNAi2 is ineffective, highlighting the need to test multiple RNAi lines.

### Depletion of Grip91^GCP3^ leads to the loss of γ-tubulin-GFP from the somatic Golgi

After having identified Grip91^GCP3^-RNAi as the most effective γ-TuRC RNAi line, we switched back to larval da neurons in order to assess the effect of Grip91 depletion on endogenously-regulated γ-tubulin-GFP localisation within the soma of da neurons. While we had previously reported that γ-tubulin-GFP associated with the somatic Golgi of these cells, it remained possible that this pool of γ-tubulin was not part of multi-protein γ-TuRCs. As previously observed, in larval da neurons not expressing RNAi we could clearly detect γ-tubulin-GFP at the somatic Golgi stacks of da neurons (Figure 2A-C, H) Strikingly, when we used 221-Gal4 to express Grip91^GCP3^-RNAi specifically within class I da neurons (ddaE and ddaD) the γ-tubulin-GFP signal was no longer apparent at the somatic Golgi stacks within these neurons, while it remained present in neighbouring da and ES neurons that did not express the RNAi (Figure 2D-F,I). A quantification of γ-tubulin-GFP signal at Golgi stacks of class I da neurons either expressing or not expressing Grip91^GCP3^-RNAi confirmed that depleting Grip91^GCP3^ leads to an almost complete loss of γ-tubulin-GFP from the somatic Golgi (Figure 2G). This loss of γ-tubulin-GFP from the Golgi was not due to a general loss of γ-tubulin-GFP expression or stability within the cell, because a diffuse γ-tubulin-GFP signal was readily detectable throughout the cytosol of Grip91^GCP3^-RNAi neurons (Figure 2I), appearing at higher levels than the cytosolic signal observed in control neurons (Figure 2H). Thus, depletion of the essential γ-TuRC protein Grip91^GCP3^ strongly inhibits the association of γ-tubulin at the somatic Golgi, confirming that Grip91^GCP3^-RNAi efficiently depletes γ-TuRCs and proving that the γ-tubulin-GFP associated with the somatic Golgi is part of *bona-fide* γ-TuRCs.

**Figure 2.**
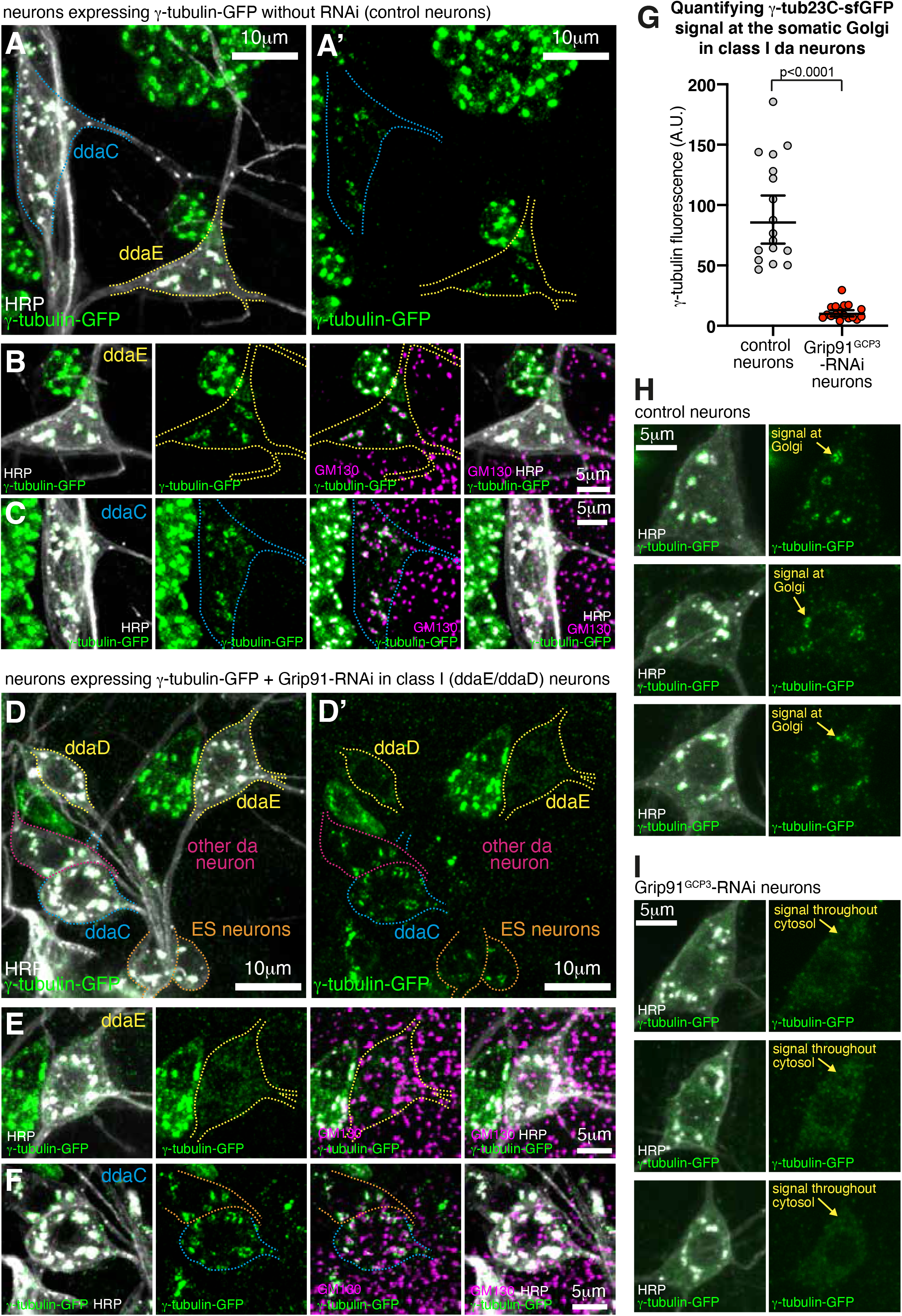
Depletion of Grip91^GCP3^ results in loss of γ-tubulin-GFP from somatic Golgi stacks. **(A-F)** Confocal images show the somas of sensory neurons within the dorsal cluster of 3^rd^ instar larva expressing endogenously-tagged γ-tubulin-GFP alone (A-C) or γ-tubulin-GFP and UAS-Grip91^GCP3^-RNAi (expressed by 221-Gal4 specifically within class I ddaD and ddaE neurons) (D-F) and immunostained for GFP (green), HRP (marking membranes including Golgi stacks, greyscale), and GM130 (marking cis-Golgi, magenta). (A, A’) and (D, D’) show a wide view of the dorsal clusters, with (A) and (D) showing the overlay of the γ-tubulin-GFP and HRP signals and (A’) and (D’) showing just the γ-tubulin-GFP signal. (B), (C), (E), and (F) display class I ddaE neurons (B,E), which express the RNAi constructs, and class IV ddaC neurons (C,F), which do not express the RNAi constructs. Note how γ-tubulin-GFP associates with HRP and GM130 in da neurons under normal conditions (A-C) but does not associate with HRP and GM130 in class I da neurons when Grip91^GCP3^-RNAi is expressed (D,E). Note also how γ-tubulin-GFP still associates with Golgi within both other da and ES neurons that do not express Grip91^GCP3^-RNAi (D,F). Scale bars are indicated within the images. (**G**) Graph shows fluorescence intensity measurements of somatic Golgi stacks within either class I da neurons expressing γ-tubulin-GFP alone (grey, N = 218 Golgi stacks from 17 neurons) or class I da neurons expressing both γ-tubulin-GFP and Grip91^GCP3^-RNAi (red, N = 249 Golgi stacks from 17 neurons). P<0.0001, unpaired t-test on log10 values. (**H, I**) Images showing examples of class I ddaE neuron soma from 3^rd^ instar larva expressing endogenously-tagged γ-tubulin-GFP alone (H, control neurons) or γ-tubulin-GFP and UAS-Grip91^GCP3^-RNAi (I, Grip91^GCP3^-RNAi neurons) and immunostained for GFP (green) and HRP (greyscale). Note how the γ-tubulin-GFP signal is predominantly concentrated at Golgi stacks in control neurons but is dispersed throughout the cytosol in Grip91^GCP3^-RNAi neurons.

### γ-TuRCs are required for asymmetric microtubule nucleation from the somatic Golgi

Having found a way to efficiently deplete γ-TuRCs from the somatic Golgi stacks of da neurons, we next wanted to test what effect this might have on microtubule growth from the Golgi. To do this we expressed either a control RNAi (γ-tubulin37C-RNAi) or Grip91^GCP3^-RNAi in conjunction with markers of microtubule growth (EB1-GFP) and Golgi stacks (ManII-mCherry) within class I da neurons. As we had observed previously (Mukherjee et al., 2020), EB1-GFP comets originated from the somatic Golgi stacks within control RNAi neurons (Figure 3A; Movie 1). Somewhat surprisingly, we found that EB1-GFP comets could still originate from Golgi stacks in Grip91^GCP3^-RNAi neurons (Figure 3B; Movie 2), and there was no significant difference in the frequency of comets originating from Golgi stacks between control-RNAi and Grip91^GCP3^-RNAi neurons (Figure 3C, unpaired t-test, p=0.64; Figure 3D, Chi-squared test for trend = 0.1131, p=0.74). Thus, γ-TuRCs do not appear to be required *per se* for microtubules to originate from the somatic Golgi, possibly because other microtubule associated proteins that promote microtubule nucleation could be present.

**Figure 3.**
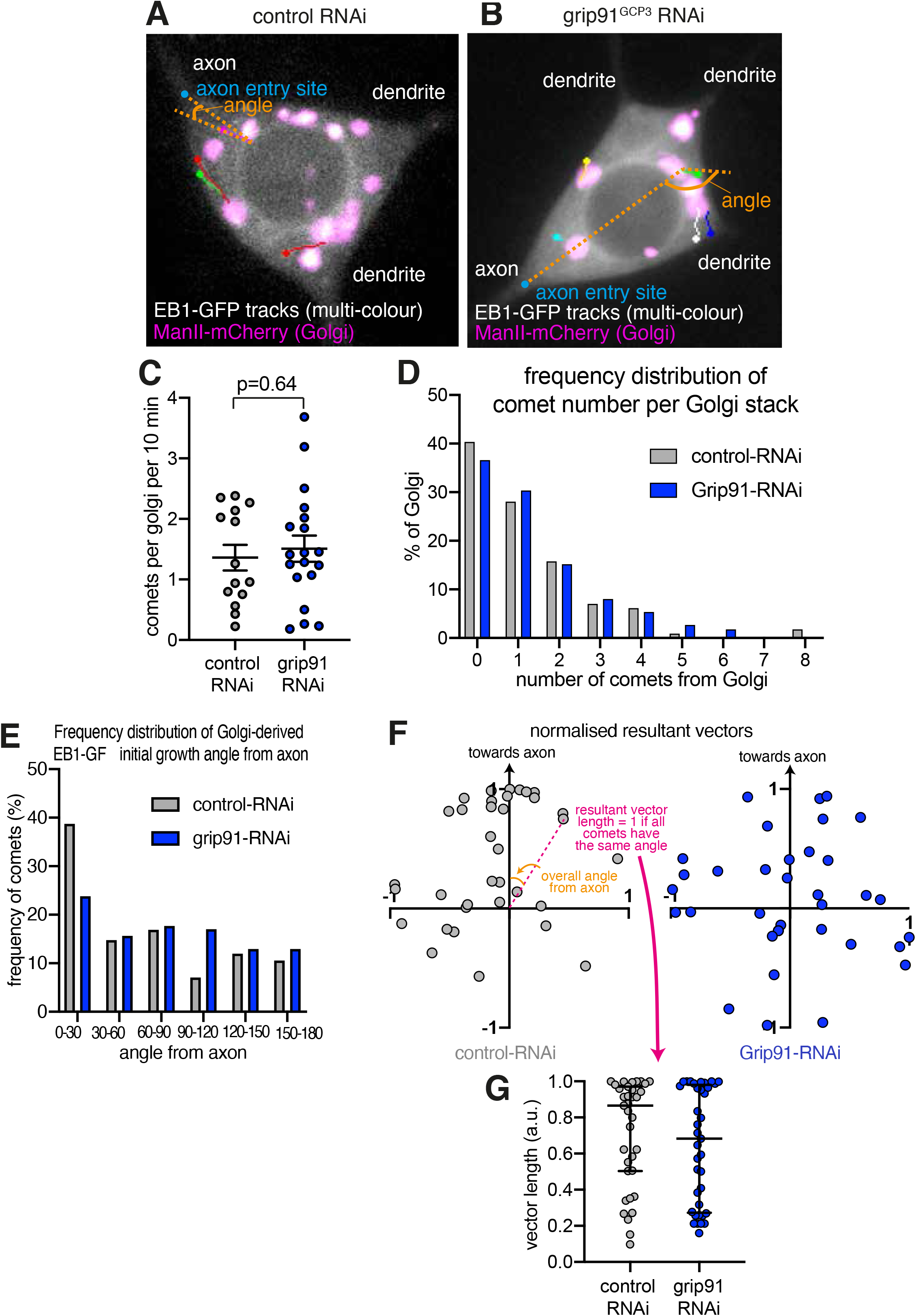
Depletion of γ-TuRCs randomises the direction of microtubule growth from somatic Golgi stacks. **(A,B)** Widefield fluorescent images from movies showing the somas of either control-RNAi (A) or Grip91^GCP3^-RNAi (B) class I ddaE neurons expressing EB1-GFP (greyscale) and ManII-mCherry (magenta). Image-J-assigned multi-colour tracks of EB1-GFP comets were drawn over each image (filled circle = last location of comet). Dotted lines show examples of angle measurements of comet initial growth relative to the axon entry point (blue dot). (**C**) Graph shows the average frequency of comets that originate from Golgi stacks (comets per Golgi stack per 10 minutes) from either control-RNAi (grey) or Grip91^GCP3^-RNAi (blue) neurons. Each point on the graph represents a single neuron, where each neuron contains multiple Golgi stacks. P=0.64, unpaired t-test. (**D**) Graph shows a frequency distribution of the number of comets that emerge from individual Golgi stacks within either control-RNAi (grey) or Grip91^GCP3^-RNAi (blue) neurons. Chi-squared test for trend = 0.1131, p=0.74. (**E**) Graph shows a frequency distribution of the initial growth angles (before turning) relative to the axon entry site (as indicated in (A,B)) of EB1-GFP comets that emerged from Golgi stacks in either control-RNAi (grey) or Grip91^GCP3^-RNAi (blue) neurons. Negative angles were made positive so as not to distinguish between comets growing to the left or right of the axon. Chi-squared=11.09, DF=5, p=0.0495. (**F**) Scatter graph shows the position of the normalised resultant comet vectors (see methods) for each Golgi stack within either control-RNAi (grey) or Grip91^GCP3^-RNAi (blue) neurons. The angle between the positive Y axis and a line drawn between a given point and the origin represents the average angle from the axon. The distance from the origin represents how similar the angles were (distance of 1 means that all comets from a given Golgi stack grew in the same direction). (**G**) Graph shows the lengths of the resultant vectors for each Golgi stack within either control-RNAi (grey) or Grip91^GCP3^-RNAi (blue) neurons. The median and interquartile range is indicated. For the data in (C-G), 114 and 112 Golgi stacks from 14 and 19 neurons were analysed for control-RNAi and Grip91^GCP3^-RNAi conditions, respectively. A total of 141 and 146 comets from control-RNAi and Grip91^GCP3^-RNAi neurons were measured, respectively.

While γ-TuRCs appear to be dispensable for microtubule nucleation from the somatic Golgi in *Drosophila* da neurons, we considered that they may still be required for the observed asymmetry in the initial microtubule growth from the Golgi i.e. γ-TuRCs may help ensure that microtubules originating from the Golgi grow preferentially towards the axon, and by extension help regulate overall microtubule polarity. In human epithelial cells, it has been proposed that a 2-step system could generate the observed asymmetric microtubule growth from the Golgi, where γ-TuRCs located at the cis-Golgi nucleate microtubules whose growth is then stabilised and promoted by the plusend-associated protein CLASP located at the trans-Golgi (Efimov et al., 2007; Vinogradova et al., 2009). To test whether γ-TuRCs are required for asymmetric microtubule nucleation from the somatic Golgi stacks in *Drosophila* da neurons we measured the initial angle of EB1-GFP comet growth relative to the axon entry site within either control-RNAi or Grip91^GCP3^-RNAi neurons. As we had found previously (Mukherjee et al., 2020), there was a strong preference for EB1-GFP comets to initially grow towards the axon in control-RNAi neurons, with ~39% of comets growing with an angle of <30° from the axon (Figure 3E). In contrast, in Grip91^GCP3^-RNAi neurons the distribution of angles across the 180° range was more evenly spread, with only ~24% of comets growing with an angle of <30° from the axon (Figure 3E). These distributions were significantly different (Chi-square=11.09, DF=5, p=0.0495). Moreover, the observed frequencies of angles from control RNAi neurons were significantly different from “expected” randomly distributed angles (there would be on average 16.7% of comets in each angle category if the angle of growth were random) (Chi-square=52.49, DF=5, p<0.0001), but this was not true in Grip91 RNAi neurons (Chi-square=7.12, DF=5, p=0.21). Thus, microtubules that originate from Golgi stacks lacking γ-TuRCs grow in a seemingly random direction relative to the axon.

We also tested the consistency of the direction of repeatedly-generated comets from each individual Golgi stack. We had previously used a resultant vector analysis to show that repeated comets emerging from a given Golgi stack tend to grow in a similar direction to each other in control-RNAi cells (Mukherjee et al., 2020). In this analysis, the distance of a vector from 0,0 (d) is equal to 1 if all comets from that particular Golgi stack travel in exactly the same direction. Meanwhile, the angle of the vector from the positive Y-axis represents the overall (or “average”) comet angle from the axon. We therefore generated normalised resultant vectors for Golgi stacks from either control RNAi or Grip91^GCP3^-RNAi cells that had produced two or more comets and plotted them on an X-Y plot (Figure 3F). As we had found previously for control-RNAi neurons, a large proportion of vectors generated from control-RNAi neurons (28/35, 80%) were plotted in the upper quadrants of the scatterplot (Figure 3F, left panel), further showing preferential growth towards the axon. There was also a bias for these vectors to be relatively long, with 20/35 (~57%) vectors having a length d > 0.8 (Figure 3G), meaning that over half of the Golgi stacks analysed generated comets they grew in a very similar direction to each other. Both of these measures were reduced in Grip91^GCP3^-RNAi neurons, where only 19/37 (~51.4%) vectors were plotted in the upper quadrants of the scatterplot (Figure 3F, right panel) and only 15/37 (40.5%) had d values > 0.8 (Figure 3G). Thus, depletion of γ-TuRCs not only randomises the direction of microtubule growth from Golgi stacks, but it also reduces the directional consistency with which Golgi-derived microtubules grow relative to one another. These results suggest that the asymmetric localisation of γ-TuRCs to the cis-face of the somatic Golgi stacks within *Drosophila* da neurons plays a decisive role in the asymmetric growth of microtubules from the Golgi.

### Turning of microtubule plus ends correlates with the position of microtubule bundles within the soma of da neurons

Along with asymmetric nucleation of microtubules from the somatic Golgi towards axons, we also recently demonstrated that the growing ends of microtubules frequently turn within the soma of *Drosophila* da neurons (Mukherjee et al., 2020). Microtubules that originate in the soma can grow into the neurites. Given the opposite microtubule polarities within axons (predominantly plus-end-out) and dendrites (predominantly minus-ends-out) it would make sense that the entry of growing plus ends into the neurites would be regulated. Indeed, we found that turning events within the soma tended to lead microtubules towards the axon and away from dendrites. Moreover, we observed that any microtubules that did happen to approach the entrance to a dendrite did not frequently enter, unlike at the axon entry site. Both microtubule turning and restricted entry into dendrites was dependent on the plus-end-directed motor Kinesin-2 (Mukherjee et al., 2020), which has been proposed to associate with, and guide, microtubule plus ends along adjacent microtubules (Chen et al., 2014; Doodhi et al., 2014; Mattie et al., 2010). We had therefore proposed that plus-end-associated Kinesin-2 guides growing microtubules along an asymmetric microtubule network generated by asymmetric nucleation from the somatic Golgi, and that plus-end-associated Kinesin-2 also limits microtubule plus end entry into dendrites when it engages with dendritic microtubules of opposite polarity (Mukherjee et al., 2020). Nevertheless, we had yet to directly show microtubule plus ends turning and growing along other microtubules within the soma of da neurons and it remained possible that the effects of Kinesin-2 depletion were indirect.

Addressing these concerns in turn, we first performed experiments to simultaneously visualise the growing ends of microtubules and microtubule bundles by co-expressing EB1-GFP and an RFP tagged microtubule marker. The microtubule binding protein mCherry-Jupiter has previously been used to label microtubule bundles within branchpoints of class I da neurons (Weiner et al., 2016), but we found that it labelled microtubules poorly within the soma and readily formed large aggregates (Movie 3), presumably due to over-expression and possibly due to a build-up of mCherry within lysosomes (Katayama et al., 2008). We therefore tested a human mCherry-α-tubulin construct that has previously been used as a microtubule marker in flies (Herzmann et al., 2018; Kanamori et al., 2015). This labelled microtubules within the soma better than mCherry-Jupiter, despite still forming aggregates (Figure 4A’-D’). The microtubule bundles remained stationary during filming but photobleached rapidly. We therefore overlayed hand-drawn lines that corresponded to the position of the microtubule bundles at the start of the movie and used them to analyse the growth trajectories of EB1-GFP comets throughout the movie. For better visualisation of the EB1-GFP comets, we tracked them using the manual tracking plugin in Fiji to generate coloured lines that represented the entire path of the comet (with a dot to indicate the final position of the comet). We overlayed these complete comet tracks independent of the time they occurred to visualise multiple comet tracks within a single image (Figure 4A-D).

**Figure 4.**
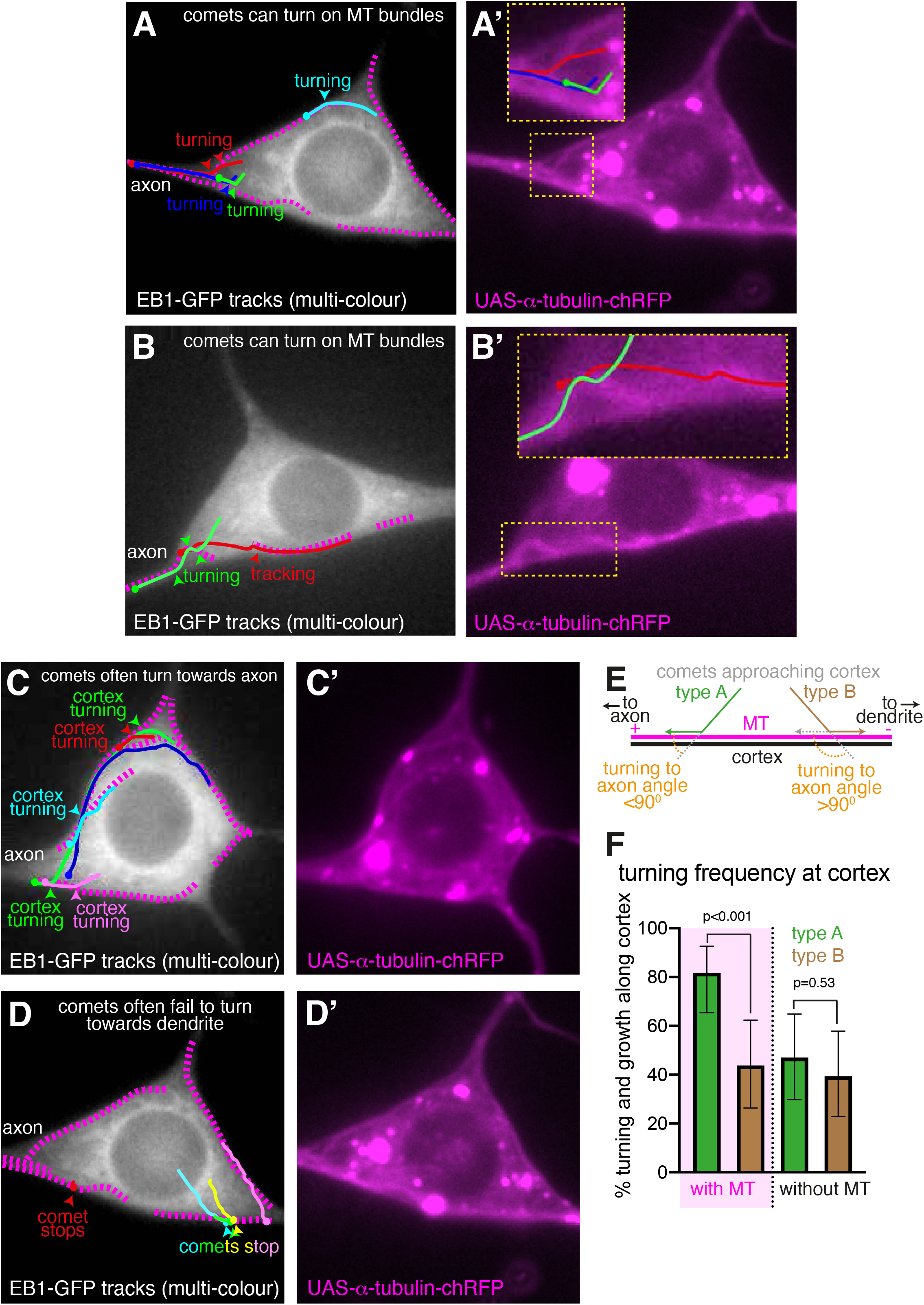
EB1-GFP comets turn along microtubule bundles within the soma of class I da neurons. **(A-D)** Widefield fluorescent images from movies showing the somas of class I da neurons expressing EB1-GFP (greyscale) (A,B,C,D) and α-tubulin-chRFP (magenta) (A’,B’,C’,D’). Image-J-assigned multi-colour tracks of EB1-GFP comets were drawn over each image (filled circle = last location of comet) in (A-D). The tracks are drawn independent of the time in which they were present in the movie, in order to allow visualisation of multiple comets in a single image. Dashed magenta lines drawn in (A-D) represent the position of microtubule bundles observed in (A’-D’) and allow the position of EB1-GFP comet tracks to be more easily compared to the position of microtubule bundles. (A) and (B) show examples of comets turning at, or following along, microtubule bundles located internally within the cell, while (C) and (D) show examples of comets that either turn (C) or do not turn (D) in peripheral regions of the cell that are associated with a microtubule bundle. Note how in (A) the green and dark blue comet tracks turn sharply at a microtubule bundle that extends out from the axon. The red comet track turns at a different bundle that bends inwards from the upper edge of the cell close to the axon; this comet then turns in the opposite direction at the other bundle extending out from the axon. The light blue comet track grows along the other end of the microtubule bundle running along the upper edge of the cell, which also bends inwards away from the edge of the cell, before turning at the cell periphery to follow the bundle towards the axon. In (B) the red comet track travels along a microtubule bundle at the lower edge of the cell before following the microtubule bundle inwards as the bundle bends away from the edge of the cell. The green comet track travels towards the axon before turning sharply one way and then the other to follow a kink in a microtubule bundle that extends out from the axon. In (C) comets turn towards the axon at cell periphery regions containing a microtubule bundle. Comets in (D) are common examples of comets disappear at the cell periphery that has an associated microtubule bundle, having grown in a direction that would have made turning towards the dendrite more favourable than turning towards the axon i.e. they are type B comets indicated in (E)). (**E**) Diagram indicating how type A and type B comets are defined. (**F**) Graph shows the frequency of type A and type B comets that turn at peripheral regions with and without microtubule bundles, as indicated. Error bars show the 95% confidence intervals. Frequencies were compared using one-way Chi squared tests.

We observed several independent examples across multiple neurons where EB1-GFP comets turned at sites that overlapped microtubule bundles (Figure 4A-D, see Figure legends for details of individual events). This included examples of comets turning along microtubule bundles that project out into the soma from the axon (Figure 4A,B) and microtubules turning along microtubules that run close to the cell periphery (Figure 4A-C). We note that not all comet turning events occur at sites with a clear microtubule bundle (for example, see the green and blue comets in Figure 4C). We believe that these turning events did also occur along microtubules but that these microtubules were not visible under our imaging conditions. Overall, we found that a number of microtubule turning events did occur very close to microtubule bundles within the soma of these neurons, consistent with our model that microtubule plus ends are guided along adjacent microtubules.

After examining several movies, it was evident that microtubule bundles often extended along the edge of the cell. In our previous paper we had not considered in our quantification turning events at the cell periphery (Mukherjee et al., 2020), as turning could be forced by collisions with the cell membrane rather than by Kinesin-2 mediated guidance along microtubules. With a microtubule marker, however, we were able to assess turning events at peripheral regions that either had or did not have a clearly associated microtubule bundle. We focussed on the peripheral regions that linked the axon entry site with a dendrite entry site (as opposed to those linking two dendrite entry sites) in order to assess whether comets turned towards the axon or towards a dendrite – our hypothesis being that these microtubule bundles would have a specific polarity with their plus ends pointing towards the axon. Indeed, when comets originated at a peripheral region their subsequent movement related to whether that peripheral region had an associated microtubule bundle or not. When comets originated at a peripheral region associated with a microtubule bundle they had a strong tendency to grow along the cell’s periphery towards the axon (30/40 comets; 75%), with only 15% of comets growing towards the dendrite; 7.5% immediately disappeared and 2.5% grew directly away from the cell periphery. In contrast, comets that emerged at a peripheral region without an obvious microtubule bundle grew with approximately equal frequency towards the axon or dendrite: 17/44 comets (38.6%) grew towards the axon, with 40.9% growing towards the dendrite. Moreover, the frequency of comets that grew directly away from the cell periphery was much higher when the comets had originated at a peripheral region without a microtubule bundle (18.2%) compared to when the comets had originated at a peripheral region with a microtubule bundle (2.5%), indicating that these microtubules help tether growing microtubules at the cell periphery.

We next assessed comets that originated within the soma and then grew into the cell periphery. We categorised these comets into type A and type B depending on the angle, relative to the axon entry site, at which they approached the cell periphery. “Type A” and “type B” comets grew into the cell periphery such that an angle of <90° would be required for them to turn along the cell periphery towards the axon or towards the dendrite, respectively i.e. type A comets could more easily turn towards the axon and type B comets could more easily turn towards the dendrite (Figure 4E). We found that, upon growing into a peripheral region that contained a microtubule bundle, ~82% of type A comets turned and continued to grow along the cell periphery towards the axon (Figure 4F), while only ~44% of type B comets turned and continued to grow along the cell periphery towards the dendrite (Chi-squared=11.77, DF=1, p=0.0006) (Figure 4F). The remaining comets in both cases stopped and disappeared. In contrast, at cell peripheral regions that did not contain a microtubule bundle only 47.1% of type A comets turned and continued to grow towards the axon, and this proportion was similar to the 39.4% of type B comets that turned and grew towards the dendrite (Chi-squared=0.395, DF=1, p=0.53) (Figure 4F). These results show that the probability of a microtubule turning when growing into the cell periphery depends upon whether there is a microtubule bundle associated with the cell periphery and on the direction with which the microtubule is growing prior to encountering the edge of the cell. The simplest way to interpret these results is that peripheral microtubule bundles tend to be polarised with their plus ends towards the axon and that microtubule plus ends are actively promoted to turn along these microtubule bundles towards the axon.

### APC depletion perturbs the behaviour of microtubule plus ends within the soma

As described above, we recently showed that depletion of Kinesin-2 reduces the frequency of comet turning, changes the frequency of comets that approach axons and dendrites, and increases the proportion of comets that enter dendrites (Mukherjee et al., 2020). If Kinesin-2 associates with and guides microtubule plus ends within the soma of da neurons, proteins involved in localising Kinesin-2 to plus ends should also be required for plus end guidance. It has been proposed that APC mediates the recruitment of Kinesin-2 to plus ends via interactions with EB1 and the regulatory subunit of Kinesin-2, Kap3 (Mattie et al., 2010). We therefore used RNAi to deplete APC from class I da neurons and monitored EB1-GFP comets that emerged within the soma, analysing the proportion of comets that turned within the soma and that approached and entered axons and dendrites (Movie S6; Figure S1). Overall, we found that depletion of APC led to similar, but less severe, phenotypes to those we had previously observed when depleting Kinesin-2 components. This was not unexpected because analysis within dendrites has previously shown that APC RNAi results in weaker phenotypes compared to RNAi targeting Kinesin-2 components, perhaps due to maternal loading of APC protein (Mattie et al., 2010). Of the 163 comets from 7 APC-RNAi neurons that grew for more than 2 μm and that did not travel along the edge of the cell, 75 displayed turning behaviour (45.4%), which is significantly lower than the 64.2% of comets that turn in control neurons (Mukherjee 2020) (p<0.001) (Figure S1). This is a relatively mild decrease compared to when depleting Kap-3 (where only 9.1% of comets turn (Mukherjee et al., 2020)), and this could be why we did not observe a significant change in the proportion of comets that approached axons (15.6% in controls (Mukherjee et al., 2020), 19.4% in APC-RNAi; p=0.12) or dendrites (7.1% in controls (Mukherjee et al., 2020) and 7.3% in APC-RNAi; p=0.95) (Figure S1). Nevertheless, there was a large increase in the proportion of comets entering dendrites. Considering only comets that reached a dendrite entry site, 27.7% enter in control-RNAi neurons (Mukherjee et al., 2020), whereas 66.7% enter the dendrite in APC RNAi neurons (p=0.001) (Figure S1). This is similar to the proportion that enter dendrites within Kap3 RNAi neurons (57.0%) and Klp64D RNAi neurons (70.0%) (Mukherjee et al., 2020). There was also a corresponding increase in the proportion of anterograde comets within proximal dendrites after APC depletion: in control neurons, only 11.1% of comets are anterograde (Mukherjee et al., 2020), while in APC RNAi neurons 30.6% are anterograde (p<0.001) (Figure S1). This change is similar to, although less severe than, that seen in Kap3-RNAi neurons, where 63.6% of proximal dendrite comets are anterograde (Mukherjee et al., 2020). Thus, depletion of APC has a similar, albeit less severe, effect on the behaviour of microtubule plus ends within the soma and proximal dendrites of class I da neurons, supporting a model in which APC facilitates the ability of Kinesin-2 to guide microtubule plus ends.

## Discussion

The data reported here and in our recent paper (Mukherjee et al., 2020), suggest the following model: γ-TuRCs asymmetrically localise to the cis-Golgi of the somatic Golgi stacks within *Drosophila* da neurons and promote the asymmetric nucleation of microtubules towards the axon. This helps generate a polarised microtubule network within the soma on which other growing microtubules are guided towards the axon and away from dendrites by plus-end-associated Kinesin-2. This asymmetric guidance increases the probability that growing microtubules reach the axon rather than a dendrite. In addition, microtubules that do attempt to grow into a dendrite are restricted from doing so via the action of plus-end-associated Kinesin-2, which we propose engages with dendritic microtubules of opposite polarity, generating a backward stalling force on the microtubule plus end as it tries to grow into the dendrite. We propose that, collectively, asymmetric nucleation and guidance and restricted entry into dendrites contributes to plus-end-out microtubule polarity within axons and helps maintain minus-end-out microtubule polarity within dendrites.

While we had previously shown the association of endogenously-tagged γ-tubulin-GFP with the cis-face of the Golgi (Mukherjee et al., 2020), we had yet to show that this pool of γ-tubulin formed part of multi-protein γ-tubulin ring complexes that promoted microtubule nucleation. We now show that depletion of the essential γ-TuRC protein Grip91^GCP3^, which is one of the proteins that holds γ-tubulin in place within the γ-TuRC, results in the loss of γ-tubulin-GFP from the Golgi, with the γ-tubulin-GFP signal still apparent in the cytosol. This is fully consistent with the conclusion that γ-tubulin-GFP exists as part of γ-TuRCs at the somatic Golgi of *Drosophila* da neurons.

In addition to Grip91^GCP3^ depletion resulting in the loss of γ-tubulin-GFP from the Golgi, we also show that there is a functional effect on microtubule growth from the Golgi. Strikingly, while depletion of Grip91^GCP3^ does not prevent microtubule nucleation from the Golgi, it does randomise the direction in which microtubules initially grow from the Golgi (discussed further below). The ability of microtubules to originate from the Golgi in the apparent absence of γ-TuRCs was somewhat surprising to us, but there is accumulating evidence that microtubules can be nucleated independently of γ-TuRCs both *in vitro* and *in vivo* (Coquand et al., 2021; Flor-Parra et al., 2018; Hannak et al., 2002; Hussmann et al., 2016; King et al., 2020; Llamazares et al., 1999; Nashchekin et al., 2016; Popov et al., 2002; Roostalu et al., 2015; Sampaio et al., 2001; Slep and Vale, 2007; Tanaka et al., 2012; Thawani et al., 2018; Tsuchiya and Goshima, 2021; Wang et al., 2015; Wieczorek et al., 2015; Woodruff et al., 2017; Zheng et al., 2020). Moreover, there is clear evidence that microtubule growth originates from Golgi outposts within the dendrites of *Drosophila* da neurons (Ori-McKenney et al., 2012; Yalgin et al., 2015; Yang and Wildonger, 2020; Zhou et al., 2014) even though the vast majority of these Golgi outposts do not associate with γ-tubulin (Mukherjee et al., 2020). Consistent with γ-TuRC-independent microtubule nucleation occurring at Golgi outposts, the ability of mispositioned Golgi outposts to affect microtubule polarity within axons is not suppressed by the depletion of γ-TuRC proteins (Arthur et al., 2015; Yang and Wildonger, 2020). Thus, if Golgi outposts can nucleate microtubules independently of γ-TuRCs it would stand to reason that the somatic Golgi is also capable of this. While we cannot rule out that a low level of γ-TuRCs could remain at the Golgi after Grip91^GCP3^-RNAi and would be sufficient to nucleate microtubules at a similar frequency to controls, we favour a hypothesis in which other microtubule associated proteins present at the Golgi can promote microtubule nucleation independently of γ-TuRCs.

The best candidates for proteins that promote γ-TuRC-independent microtubule nucleation are the homologues of CAMSAP, ch-TOG, and CLASP, which can all bind microtubules directly. CAMSAP proteins specifically bind and regulate microtubule minus ends (Atherton et al., 2017; Hendershott and Vale, 2014), while ch-TOG and CLASP bind and regulate plus ends (Slep, 2009). Both CAMSAP and CLASP proteins have already been shown to function in microtubule organisation at the somatic Golgi of mammalian cells (Efimov et al., 2007; Wu et al., 2016). While ch-TOG or its homologues have not been reported to localise to Golgi structures, TOG domain proteins can clearly promote microtubule nucleation in the absence of γ-TuRCs *in vitro* (Flor-Parra et al., 2018; Hussmann et al., 2016; King et al., 2020; Popov et al., 2002; Roostalu et al., 2015; Slep and Vale, 2007; Thawani et al., 2018; Wieczorek et al., 2015; Woodruff et al., 2017) and *in vivo* (Hannak et al., 2002; Tsuchiya and Goshima, 2021; Zheng et al., 2020). CLASP can also to promote microtubule nucleation in human colon cancer cells after γ-TuRC depletion (Tsuchiya and Goshima, 2021). Nevertheless, whether the *Drosophila* homologues of CAMSAP, ch-TOG and CLASP proteins localise to the somatic Golgi and function in microtubule nucleation in da neurons remains unknown.

Although we found no apparent effect on microtubule nucleation, depletion of γ-TuRCs clearly randomised the direction of initial microtubule growth from the somatic Golgi. It is possible that γ-TuRCs function in conjunction with the plus-end-binding proteins CLASP or Msps to ensure that microtubules grow predominantly towards the axon after nucleation. A model has been proposed in mammalian cells in which trans-Golgi associated CLASP stabilises and promotes the growth of microtubules nucleated by γ-TuRCs at the cis-Golgi, thereby generating an asymmetry in microtubule nucleation (Vinogradova et al., 2009). This could in theory also occur within *Drosophila* da neurons with the possibility that CLASP, or other microtubule binding proteins, could themselves stimulate microtubule assembly after depletion of γ-TuRCs, with the resulting microtubules growing in any orientation. This, however, remains to be tested.

In addition to showing a role for γ-TuRCs in asymmetric microtubule nucleation from the Golgi, we have provided further evidence that growing microtubule plus ends are guided along a polarised microtubule network within the soma to help regulate microtubule polarity within neurites. We show that the plus ends of growing microtubules turn at locations in the soma where microtubule bundles are present, strongly suggesting that turning occurs directly on microtubules. We also show that turning at the cell periphery is influenced by the presence of peripheral microtubule bundles and on the direction in which the microtubule is growing when it encounters the edge of the cell. From these results we conclude that the peripheral microtubule bundles are polarised with respect to axons and dendrites and that they play an instructive role in the behaviour of growing microtubules when they encounter the edge of the cell.

In our previous paper we had shown that microtubule guidance within the neuronal soma was dependent on Kinesin-2 and proposed that it was the association of Kinesin-2 with microtubule plus ends that was necessary for their guidance (Mukherjee et al., 2020). Here we show that depletion of APC, the protein proposed to link Kinesin-2 to the plus ends of growing microtubules, results in similar phenotypes to depletion of Kinesin-2 i.e. it reduces the proportion of microtubule plus ends that turn within the soma and increases the proportion that enter dendrites, resulting in an increase in plus-end-out microtubules within the proximal dendrites. This supports the model that APC links Kinesin-2 to microtubule plus ends to allow Kinesin-2 to mediate plus end guidance.

In summary, γ-TuRCs asymmetrically localised at the somatic Golgi of *Drosophila* class I da neurons play an important role in asymmetric microtubule nucleation. This asymmetric nucleation appears to help generate an asymmetric microtubule network within the soma on which growing microtubule plus ends are guided by Kinesin-2 towards axons and away from dendrites. Our data is consistent with a model in which Kinesin-2 associates with microtubule plus via an interaction with APC. Collectively, these processes help establish and maintain microtubule polarity within neurites, by increasing the plus-end-out microtubule pool within axons and restricting the number of plus ends that grow into dendrites. It will be interesting in future to see whether such tight regulation of somatic microtubules is of general importance across neuronal types and across species.

## Acknowledgements

This work was supported by a Wellcome Trust and Royal Society Sir Henry Dale Fellowship (105653/Z/14/Z), an Isaac Newton Trust Research grant (18.23(p)), and an IdEx Université de Paris ANR-18-IDEX-0001 awarded to PTC. We thank Caroline Fabre for help with tracking of EB1-GFP comets. We thank members of the Conduit lab for their invaluable input and critical reading of the manuscript. We thank Clemens Cabernard and Yuh-Nung Jan for fly lines that were used in this study. The work benefited from use of the Imaging Facility, Department of Zoology, supported by Matt Wayland and a Sir Isaac Newton Trust Research Grant (18.07ii(c)). For the purpose of Open Access, the author has applied a CC BY public copyright license to any Author Accepted Manuscript version arising from this submission. The authors declare no financial or non-financial competing interests.

## Supplementary Figure and Movie Legends

**Figure S1.**
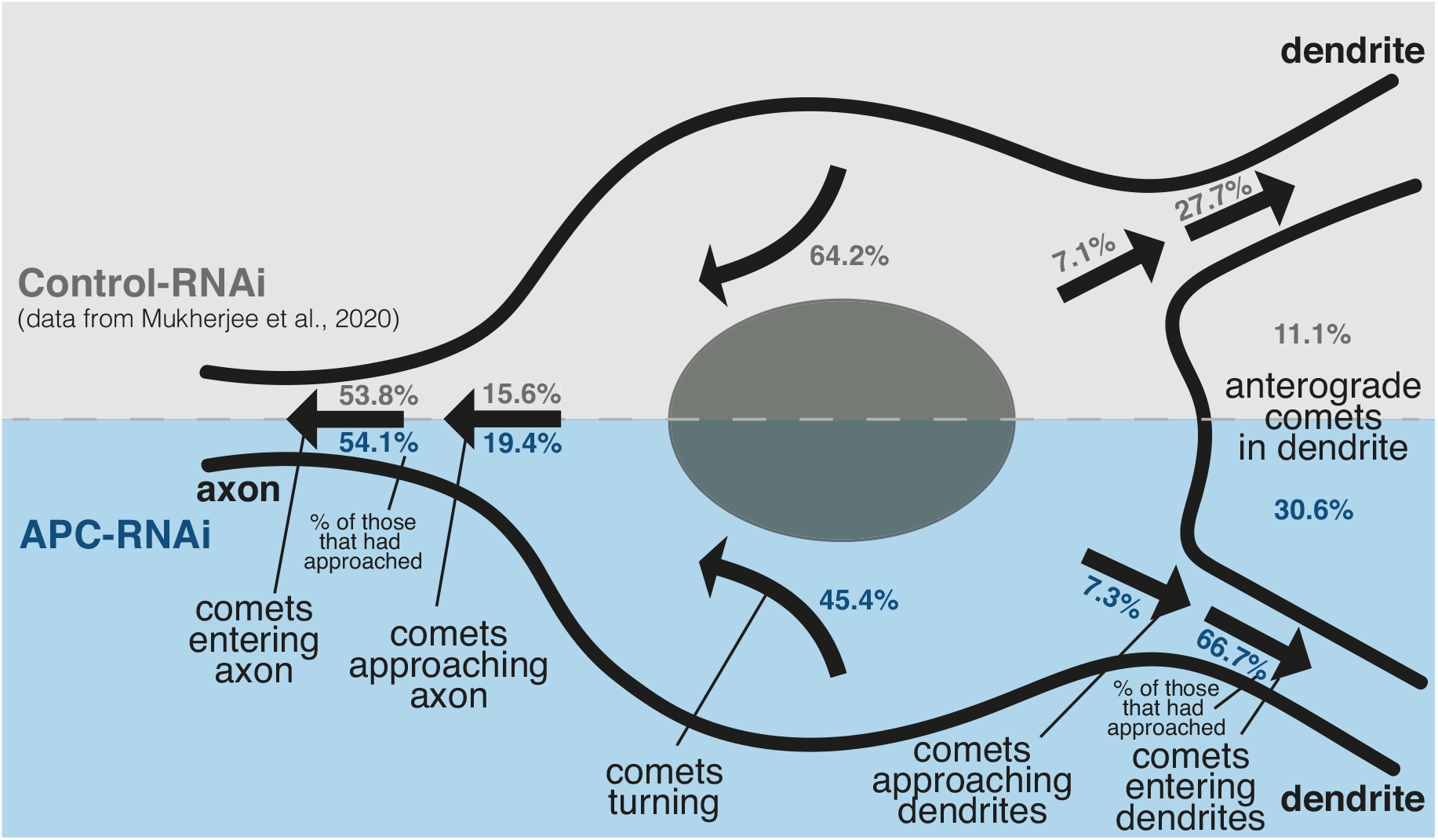
Summary of EB1-GFP comet behaviour in wildtype and APC RNAi knockdown neurons. Cartoon summarising the behaviour of EB1-GFP comets within the soma of control-RNAi (upper section, data taken from Mukherjee et al., 2020) and APC-RNAi (lower section, this study) neurons. Proportions of comets that grew for more than 2μm and that turned within the soma (n= 257 and 163 comets from 13 and 7 control-RNAi and APC-RNAi neurons, respectively), that approached axons or dendrites (n=666 and 371 comets from 13 and 7 control-RNAi and APC-RNAi neurons, respectively), that enter axons (n=104 and 72 comets from 13 and 7 control-RNAi and APC-RNAi neurons, respectively) or dendrites (n=47 and 27 comets from 13 and 7 control-RNAi and APC-RNAi neurons, respectively) once they have approached, and of anterograde comets within proximal dendrites (n=252 and 147 comets from 13 and 7 control-RNAi and APC-RNAi neurons, respectively) are indicated in black (control-RNAi) and dark blue (APC-RNAi).

**Movie S1 – related to Figure 3.**

**Microtubules originating from the somatic Golgi stacks grow preferentially towards the axon in control neurons.** The Movie was taken using single-Z timelapse epifluorescence microscopy and shows a class I da neuron expressing UAS-Dicer2, UAS-γ-tubulin37C-RNAi, UAS-EB1-GFP and UAS-ManII-mCherry driven by 221-Gal4. ImageJ-generated EB1-GFP comet line tracks are shown only for the microtubule growth events that originate at the somatic Golgi stacks. The tracks are multi-coloured, as is default for ImageJ, and the colours have no reference to comet type. The axon and dendrites are labelled. Comets can be seen emerging from Golgi stacks and growing towards the axon; this can occur repeatedly from the same stack (see stack in left centre of soma). Images were collected every 3s and the Movie plays at 11 frames/second.

**Movie S2 – related to Figure 3.**

**Microtubules originating from the somatic Golgi stacks grow in random directions with respect to the axon in Grip91^GCP3^-RNAi neurons.** The Movie was taken using single-Z time-lapse epifluorescence microscopy and shows a class I da neuron expressing UAS-Dicer2, UAS-Grip91^GCP3^-RNAi, UAS-EB1-GFP and UAS-ManII-mCherry driven by 221-Gal4. ImageJ-generated EB1-GFP comet line tracks are shown only for the microtubule growth events that originate at the somatic Golgi stacks. The tracks are multi-coloured, as is default for ImageJ, and the colours have no reference to comet type. The axon and dendrites are labelled. Comets can be seen emerging from Golgi stacks in various directions with respect to the axon. Images were collected every 3s and the Movie plays at 11 frames/second.

**Movie S3 – related to Figure 4.**

**221-gal4>UAS-mCherry-Jupiter is a poor marker of microtubules within the soma of class I da neurons.** The Movie was taken using single-Z time-lapse epifluorescence microscopy and shows a class I da neuron expressing UAS-EB1-GFP and UAS-mCherry-Jupiter driven by 221-Gal4 (only the UAS-mCherry-Jupiter signal is shown). Note how mCherry-Jupiter forms large aggregates within the soma and that microtubule bundles are not readily visible. Images were collected every 3s and the Movie plays at 7 frames/second.

**Movie S4 – related to Figure 4.**

**EB1-GFP comets turn towards the axon within the soma of class I da neurons.** The Movie was taken using single-Z time-lapse epifluorescence microscopy and shows a class I da neuron expressing UAS-EB1-GFP (greyscale) and UAS-mCherry-α-tubulin by 221-Gal4 (only the UAS-EB1-GFP signal is shown). Magenta arrowheads highlight selected comets that turn towards the axon close to the axon entry site. These comets are tuning on microtubules bundles, as indicated in Figure 4A. Note how the comet that starts at timepoint 198s turns first one way and then the other along separate microtubule bundles (see Figure 4A). The comet that starts at timepoint 240s is not a turning comet but it splits into two comets at timepoint 291s, which grow along two different microtubule bundles extending out from the axon (see Figure 4A’). Yellow arrowheads indicate type B comets (i.e. those growing more towards a dendrite than the axon, see Figure 4E) that fail to turn at the cell periphery, which contains a microtubule bundle (see Figure 4D). Images were collected every 3s and the Movie plays at 7 frames/second.

**Movie S5 – related to Figure 4.**

**EB1-GFP comets turn along microtubule bundles within the soma of class I da neurons.** The Movie was taken using single-Z time-lapse epifluorescence microscopy and shows a class I da neuron expressing UAS-EB1-GFP (greyscale) and UAS-mCherry-α-tubulin by 221-Gal4 (magenta). Only a selection of the UAS-mCherry-α-tubulin signal is shown and is labelled ‘microtubules’. Magenta arrowheads highlight selected comets that appear to interact with microtubule bundles. Note how the 2 comets that grow down towards the axon pause when they encounter the bent microtubule bundle that protrudes from the axon and then change direction to grow along the bend in the microtubule. Note also how the comet growing along the lower edge of the cell grows inwards at the point where the microtubule bundle turns inwards. Images were collected every 3s and the Movie plays at 7 frames/second.

**Movie S6 – related to Figure 4 and to Figure S1.**

**Depletion of APC perturbs the behaviour of microtubule plus ends within the soma of da neurons.** The Movie was taken using single-Z time-lapse epifluorescence microscopy and shows a class I da neuron expressing UAS-Dicer2, UAS-APC-RNAi and UAS-EB1-GFP driven by 221-Gal4. The left panel shows the original movie and the right panel shows the original movie with overlayed imageJ-generated multicoloured EB1-GFP comet line tracks dots. The different colours are set by default for ImageJ and have no reference to comet type. The axon and dendrites are labelled. In this particular movie, only 38.6% of comets that grow for more than 2μm turn within the soma. 14.8% and 12.3% of comets approach the axon or a dendrite, respectively. Of these comets, 50% enter the axon and 90% enter a dendrite. Images were collected every 5s and the Movie plays at 7 frames/second.

## Materials and methods

### Contact for Reagent and Resource Sharing

Further information and requests for resources and reagents should be directed to and will be fulfilled by the Lead Contact, Paul Conduit.

### Experimental Model and Subject Details

All fly strains were maintained at 18 or 25°C on Iberian fly food made from dry active yeast, agar, and organic pasta flour, supplemented with nipagin, propionic acid, pen/strep and food colouring.

### Methods

#### *Drosophila melanogaster* stocks

The following fluorescent alleles were used in this study: ppk-CD4-tdGFP (BL 35842), γ-tubulin23C-sfGFP (Tovey et al., 2018), γ-tubulin23c-eGFP (Mukherjee et al., 2020), UAS-mCherry-Jupiter (gift from Clemens Cabernard), UAS-ChRFP-Tubulin (BL 25774), UAS-EB1-GFP (BL 35512), UAS-ManII-mCherry (gift from Yuh-Nung Jan). The following Gal4 and RNAi lines were used in this study: ppk-Gal4 (BL 32078), 221-Gal4 (BL 26259), UAS-Grip91^GCP3^-RNAi1 (VDRC 104667), UAS-Grip 91^GCP3^-RNAi2 (BL 31201), UAS-Grip84^GCP2^-RNAi1 (VDRC 105640), UAS-Grip84^GCP2^-RNAi2 (BL 33548), γ-tubulin37C-RNAi (BL32513), γ-tubulin23C-RNAi1 (BL 31204), γ-tubulin23C-RNAi2 (VDRC 19130), UAS-Dicer 2 (BL 24646), UAS-Apc-RNAi (VDRC 51468).

For examining the morphology of pupal class IV v’ada neurons we used flies expressing one copy of ppk-CD4-tdGFP and containing one copy of UAS-Dicer2 and one copy of the appropriate UAS-driven RNAi line expressed under the control of one copy of ppk-Gal4. For examining the endogenous localisation of γ-tubulin23C we used flies expressing γ-tubulin23C-sfGFP and γ-tubulin23C-eGFP i.e. 2 copies of γ-tubulin23C-GFP. For examining the localisation of γ-tubulin23C during Grip91^GCP3^ depletion we used flies expressing two copies of γ-tubulin23C-(sf/e)GFP (as above) in combination with one copy of UAS-Grip91^GCP3^ RNAi and UAS-Dicer2 expressed under the control of one copy of 221-Gal4. For examining microtubule dynamics in relation to the somatic Golgi we used flies expressing one copy of UAS-ManII-mCherry, one copy of UAS-EB1-GFP, one copy of UAS-Dicer 2, and either one copy of UAS-γ-tubulin37C RNAi (control) or one copy of UAS-Grip91^GCP3^ RNAi, using one copy of 221-Gal4. For examining microtubule dynamics in relation to stable microtubules we used flies with one copy of UAS-EB1-GFP and one copy of UAS-mCherry-Jupiter or UAS-ChRFP-Tubulin expressed under the control of one copy of 221-Gal4. For examining microtubule dynamics during APC depletion we used flies with one copy of UAS-EB1-GFP, one copy of UAS-Dicer 2 and one copy of UAS-APC-RNAi expressed under the control of one copy of 221-Gal4.

#### Antibodies

The following primary antibodies were used: anti-GFP mouse monoclonal at 1:250 (Roche, 11814460001), Alexa-647 conjugated HRP polyclonal at 1:500 (Jackson), and anti-GM130 rabbit polyclonal at 1:300 (Abcam). The following secondary antibodies were used: Alexa-488 anti-Mouse at 1:500 (ThermoFisher), Alexa-547 anti-Rabbit at 1:500 (ThermoFisher).

#### Fixed and live cell imaging

Imaging of pupal neurons and of γ-tubulin-GFP within larval neurons was carried out on an Olympus FV3000 scanning inverted confocal system run by FV-OSR software using a 60X 1.4NA silicone immersion lens (UPLSAPO60xSilicone). For imaging class IV v’ada neurons within pupae, white pre-pupae (0h APF) were collected and placed in a sealed petri dish with some moistened tissue paper and aged for the required duration in a 25°C incubator. The staged pupae were stuck on double-sided sticky tape and removed from their pupal casing. The pupae were mounted on a glass slide with a small amount of halocarbon oil and a coverglass was placed gently on top of the pupae using dental wax feet as spacers. The mounted pupae were imaged immediately. For live larval samples, wandering L3 larvae were placed in a drop of glycerol and flattened between a slide and a 22X22mm coverslip, held in place by double-sided sticky tape, and imaged immediately. Imaging of EB1-GFP and ManII-mCherry within da neurons was performed on a Leica DM IL LED inverted microscope controlled by μManager software and coupled to a RetigaR1 monochrome camera (QImaging) and a CoolLED pE-300 Ultra light source using a 63X 1.3NA oil objective (Leica 11506384). Single Z-plane images were acquired every 5 seconds; for EB1-GFP/ManII-mCherry or EB1-GFP/ChRFP-Tubulin or EB1-GFP/ Cherry-Jupiter, dual imaging single Z-plane images were acquired every 3 seconds. All images were processed using Fiji (ImageJ). EB1 comets were tracked using the Manual Tracking plugin in Fiji.

#### Quantification and Statistical Analysis

Data was processed in Microsoft Excel. Statistical analysis and graph production were performed using GraphPad Prism.

Analysis of the γ-tubulin-GFP signal at somatic Golgi stacks in control versus Grip91^GCP3^-RNAi neurons: Maximum intensity Z-plane projections were made and ROIs were drawn around individual Golgi stacks. Sum fluorescence intensities for each Golgi stack were calculated and then background corrected using overall mean cytosolic intensity measurements. The mean “corrected” intensity was calculated for individual neurons and plotted in Figure 2G. The data sets were compared using an unpaired t-test on log_10_ values, because the data sets were log-normally distributed.

Analysis of EB1-GFP comets was carried out in a similar way as previously (Mukherjee et. al., 2020). Briefly, comets that were present within the first timepoint (and thus may not represent newly growing microtubules) were excluded. The angle of initial comet growth was measured using the angle tool within ImageJ. The frequencies of comets per Golgi per 10 minutes in Figure 3C were compared using an unpaired t-test. The frequency distributions of comet number per Golgi stack in Figure 3D were compared using a chi-squared test for trend analysis. For the frequency distribution of comet angles in Figure 3E, the negative angles were made positive, so as not to distinguish between comets growing either side of the axon entry site, creating a distribution between 0° and 180°. A total of 141 and 146 comets from control-RNAi and Grip91^GCP3^-RNAi neurons were measured, respectively. The frequency distributions were compared to each other and to an expected distribution (if assuming random angles) using contingency chi-squared analyses. We used both positive and negative angles (between −180° and 180°) for the vector analysis in Figure 3F,G. Only Golgi stacks that generated 2 or more comets were analysed. Vectors for individual comets from a single Golgi stack were summed and then the value was normalised by dividing by the number of comets, such that the maximum possible length of the resultant vector would be 1, irrespective of comet number. 114 and 112 Golgi stacks from 14 and 19 neurons were analysed for control-RNAi and Grip91^GCP3^-RNAi conditions, respectively. Vector lengths could not easily be statistically compared due to their unusual distributions.

For the analysis of comet turning at peripheral cell regions with or without a microtubule bundle we analysed a total of 227 comets and 15 peripheral cell regions from 11 neurons. 10 peripheral cell regions contained a microtubule bundle, 5 did not. At peripheral cell regions with a microtubule bundle, 44 comets were type A and 32 were type B, while 40 comets originated at these peripheral cell regions; at peripheral cell regions without a microtubule bundle, 34 comets were type A and 33 were type B, while 44 originated at these peripheral cell regions.

For APC RNAi neurons, we analysed a total of 371 comets (across 7 movies) that had originated within the soma during the movies (average of 5.4 comets per minute). 272 of these comets disappeared within the soma, 72 approached the axon and 27 approached a dendrite. The proportion of these comets that entered the axon or dendrites was calculated. To assess the polarity of dynamic microtubules within the proximal dendritic regions, we included both comets that originated within the proximal dendrite and those that grew into the dendrite from the soma, and included comets that were already present at the start of the movie. We analysed a total of 147 comets from 7 movies. To assess the proportion of comets that turned within the soma, we considered only those comets that grew for more than 2μm within the soma and included comets that had grown into the soma from the neurites. In total, 163 comets were analysed for turning behaviour. The proportions of comets that turned within the soma, that approached axons or dendrites, that entered axons or dendrites after approaching, and that were anterograde within the proximal dendrite were compared to the proportions we had previously calculated for control-RNAi neurons (Mukherjee et al., 2020) using Chi-squared tests.

---

**Table S1.**

| lab code | gene | RNAi type | stock no. | 40D present |
| --- | --- | --- | --- | --- |
| R45 | g-tub37C | TRiP valium 20 | BL32513 | n/a |
| R1 | g-tub23C RNAi1 | TRiP valium 1 | BL31204 | n/a |
| R21 | g-tub23C RNAi2 | VDRC GD | v19130 | n/a |
| R33 | Grip84<sup>GCP2</sup> RNAi1 | VDRC KK | v105640 | no |
| R41 | Grip84<sup>GCP2</sup> RNAi2 | TRiP valium 20 | BL33548 | n/a |
| R34 | Grip91<sup>GCP3</sup> RNAi1 | VDRC KK | v104667 | no |
| R42 | Grip91<sup>GCP3</sup> RNAi2 | TRiP valium 1 | BL31201 | n/a |

## Notes

### Competing Interest Statement

The authors have declared no competing interest.

### Summary of Updates

Order of author names

## References

Arthur AL, Yang SZ, Abellaneda AM, Wildonger J. 2015. Dendrite arborization requires the dynein cofactor NudE. J Cell Sci 128:2191 2201. doi:10.1242/jcs.170316

Atherton J, Jiang K, Stangier MM, Luo Y, Hua S, Houben K, Hooff JJE van, Joseph A-P, Scarabelli G, Grant BJ, Roberts AJ, Topf M, Steinmetz MO, Baldus M, Moores CA, Akhmanova A. 2017. A structural model for microtubule minus-end recognition and protection by CAMSAP proteins. Nat Struct Mol Biology 24:nsmb.3483. doi:10.1038/nsmb.3483

Baas PW, Lin S. 2011. Hooks and comets: The story of microtubule polarity orientation in the neuron. Dev Neurobiol 71:403–418. doi:10.1002/dneu.20818

Castillo U del, Norkett R, Gelfand VI. 2019. Unconventional Roles of Cytoskeletal Mitotic Machinery in Neurodevelopment. Trends Cell Biol. doi:10.1016/j.tcb.2019.08.006

Chen L, Stone MC, Tao J, Rolls MM. 2012. Axon injury and stress trigger a microtubule-based neuroprotective pathway. Proc National Acad Sci 109:11842–11847. doi:10.1073/pnas.1121180109

Chen Y, Rolls MM, Hancock WO. 2014. An EB1-Kinesin Complex Is Sufficient to Steer Microtubule Growth In Vitro. Curr Biol 24:316–321. doi:10.1016/j.cub.2013.11.024

Conduit PT, Wainman A, Raff JW. 2015. Centrosome function and assembly in animal cells. Nat Rev Mol Cell Bio 16:611 624. doi:10.1038/nrm4062

Coquand L, Victoria GS, Tata A, Carpentieri JA, Brault J-B, Guimiot F, Fraisier V, Baffet AD. 2021. CAMSAPs organize an acentrosomal microtubule network from basal varicosities in radial glial cells. J Cell Biol 220:e202003151. doi:10.1083/jcb.202003151

Cunha-Ferreira I, Chazeau A, Buijs RR, Stucchi R, Will L, Pan X, Adolfs Y, Meer C van der, Wolthuis JC, Kahn OI, Schätzle P, Altelaar M, Pasterkamp RJ, Kapitein LC, Hoogenraad CC. 2018. The HAUS Complex Is a Key Regulator of Non-centrosomal Microtubule Organization during Neuronal Development. Cell Reports 24:791 800. doi:10.1016/j.celrep.2018.06.093

Doodhi H, Katrukha EA, Kapitein LC, Akhmanova A. 2014. Mechanical and Geometrical Constraints Control Kinesin-Based Microtubule Guidance. Curr Biol 24:322–328. doi:10.1016/j.cub.2014.01.005

Efimov A, Kharitonov A, Efimova N, Loncarek J, Miller PM, Andreyeva N, Gleeson P, Galjart N, Maia ARR, McLeod IX, Yates JR, Maiato H, Khodjakov A, Akhmanova A, Kaverina I. 2007. Asymmetric CLASP-dependent nucleation of noncentrosomal microtubules at the trans-Golgi network. Dev Cell 12:917 930. doi:10.1016/j.devcel.2007.04.002

Farache D, Emorine L, Haren L, Merdes A. 2018. Assembly and regulation of γ-tubulin complexes. Open Biol 8:170266. doi:10.1098/rsob.170266

Feng C, Thyagarajan P, Shorey M, Seebold DY, Weiner AT, Albertson RM, Rao KS, Sagasti A, Goetschius DJ, Rolls MM. 2019. Patronin-mediated minus end growth is required for dendritic microtubule polarity. J Cell Biology 218:2309–2328. doi:10.1083/jcb.201810155

Flor-Parra I, Iglesias-Romero AB, Chang F. 2018. The XMAP215 Ortholog Alp14 Promotes Microtubule Nucleation in Fission Yeast. Curr Biol 28:1681 1691.e4. doi:10.1016/j.cub.2018.04.008

Fu M, McAlear TS, Nguyen H, Oses-Prieto JA, Valenzuela A, Shi RD, Perrino JJ, Huang T-T, Burlingame AL, Bechstedt S, Barres BA. 2019. The Golgi Outpost Protein TPPP Nucleates Microtubules and Is Critical for Myelination. Cell 179:132–146.e14. doi:10.1016/j.cell.2019.08.025

Garbrecht J, Laos T, Holzer E, Dillinger M, Dammermann A. 2021. An acentriolar centrosome at the C. elegans ciliary base. Curr Biol 31:2418–2428.e8. doi:10.1016/j.cub.2021.03.023

Grueber WB, Jan LY, Jan Y-N. 2002. Tiling of the Drosophila epidermis by multidendritic sensory neurons. Development (Cambridge, England) 129:2867 2878.

Hannak E, Oegema K, Kirkham M, Gönczy P, Habermann B, Hyman AA. 2002. The kinetically dominant assembly pathway for centrosomal asters in Caenorhabditis elegans is γ-tubulin dependent. J Cell Biology 157:591–602. doi:10.1083/jcb.200202047

Harterink M, Edwards SL, Haan B de, Yau KW, Heuvel S van den, Kapitein LC, Miller KG, Hoogenraad CC. 2018. Local microtubule organization promotes cargo transport in C. elegans dendrites. J Cell Sci 131:jcs.223107. doi:10.1242/jcs.223107

Hendershott MC, Vale RD. 2014. Regulation of microtubule minus-end dynamics by CAMSAPs and Patronin. Proc National Acad Sci 111:5860 5865. doi:10.1073/pnas.1404133111

Herzmann S, Götzelmann I, Reekers L-F, Rumpf S. 2018. Spatial regulation of microtubule disruption during dendrite pruning in Drosophila. Development 145:dev156950. doi:10.1242/dev.156950

Hussmann F, Drummond DR, Peet DR, Martin DS, Cross RA. 2016. Alp7/TACC-Alp14/TOG generates long-lived, fast-growing MTs by an unconventional mechanism. Sci Rep-uk 6:20653. doi:10.1038/srep20653

Jimbo T, Kawasaki Y, Koyama R, Sato R, Takada S, Haraguchi K, Akiyama T. 2002. Identification of a link between the tumour suppressor APC and the kinesin superfamily. Nat Cell Biol 4:323–327. doi:10.1038/ncb779

Kamasaki T, O’Toole E, Kita S, Osumi M, Usukura J, McIntosh JR, Goshima G. 2013. Augmin-dependent microtubule nucleation at microtubule walls in the spindle. J Cell Biol 202:25–33. doi:10.1083/jcb.201304031

Kanamori T, Yoshino J, Yasunaga K, Dairyo Y, Emoto K. 2015. Local endocytosis triggers dendritic thinning and pruning in Drosophila sensory neurons. Nat Commun 6:6515. doi:10.1038/ncomms7515

Kapitein LC, Hoogenraad CC. 2015. Building the Neuronal Microtubule Cytoskeleton. Neuron 87:492 506. doi:10.1016/j.neuron.2015.05.046

Katayama H, Yamamoto A, Mizushima N, Yoshimori T, Miyawaki A. 2008. GFP-like Proteins Stably Accumulate in Lysosomes. Cell Struct Funct 33:1–12. doi:10.1247/csf.07011

King BR, Moritz M, Kim H, Agard DA, Asbury CL, Davis TN. 2020. XMAP215 and γ-tubulin additively promote microtubule nucleation in purified solutions. Mol Biol Cell mbcE20020160. doi:10.1091/mbc.e20-02-0160

Kollman JM, Merdes A, Mourey L, Agard DA. 2011. Microtubule nucleation by γ-tubulin complexes. Nat Rev Mol Cell Bio 12:709 721. doi:10.1038/nrm3209

Liang X, Kokes M, Fetter RD, Sallee MD, Moore AW, Feldman JL, Shen K. 2020. Growth cone-localized microtubule organizing center establishes microtubule orientation in dendrites. Elife 9:e56547. doi:10.7554/elife.56547

Lin S, Liu M, Mozgova OI, Yu W, Baas PW. 2012. Mitotic motors coregulate microtubule patterns in axons and dendrites. J Neurosci 32:14033 14049. doi:10.1523/jneurosci.3070-12.2012

Llamazares S, Tavosanis G, González C. 1999. Cytological characterisation of the mutant phenotypes produced during early embryogenesis by null and loss-of-function alleles of the gammaTub37C gene in Drosophila. Journal of cell science 112(Pt 5):659 667.

Lu W, Gelfand VI. 2017. Moonlighting Motors: Kinesin, Dynein, and Cell Polarity. Trends Cell Biol 27:505–514. doi:10.1016/j.tcb.2017.02.005

Lüders J. 2020. Nucleating microtubules in neurons: Challenges and solutions. Dev Neurobiol. doi:10.1002/dneu.22751

Magescas J, Eskinazi S, Tran MV, Feldman JL. 2021. Centriole-less pericentriolar material serves as a microtubule organizing center at the base of C. elegans sensory cilia. Curr Biol 31:2410–2417.e6. doi:10.1016/j.cub.2021.03.022

Mattie FJ, Stackpole MM, Stone MC, Clippard JR, Rudnick DA, Qiu Y, Tao J, Allender DL, Parmar M, Rolls MM. 2010. Directed microtubule growth, +TIPs, and kinesin-2 are required for uniform microtubule polarity in dendrites. Curr Biol 20:2169 2177. doi:10.1016/j.cub.2010.11.050

Mimori-Kiyosue Y, Shiina N, Tsukita S. 2000. Adenomatous Polyposis Coli (APC) Protein Moves along Microtubules and Concentrates at Their Growing Ends in Epithelial Cells. J Cell Biology 148:505–518. doi:10.1083/jcb.148.3.505

Mukherjee A, Brooks PS, Bernard F, Guichet A, Conduit PT. 2020. Microtubules originate asymmetrically at the somatic Golgi and are guided via Kinesin2 to maintain polarity in neurons. Elife 9:e58943. doi:10.7554/elife.58943

Nashchekin D, Fernandes AR, Johnston DS. 2016. Patronin/Shot Cortical Foci Assemble the Noncentrosomal Microtubule Array that Specifies the Drosophila Anterior-Posterior Axis. Dev Cell 38:61 72. doi:10.1016/j.devcel.2016.06.010

Näthke IS, Adams CL, Polakis P, Sellin JH, Nelson WJ. 1996. The adenomatous polyposis coli tumor suppressor protein localizes to plasma membrane sites involved in active cell migration. J Cell Biology 134:165–179. doi:10.1083/jcb.134.1.165

Nguyen MM, McCracken CJ, Milner ES, Goetschius DJ, Weiner AT, Long MK, Michael NL, Munro S, Rolls MM. 2014. γ-Tubulin controls neuronal microtubule polarity independently of Golgi outposts. Mol Biol Cell 25:2039–2050. doi:10.1091/mbc.e13-09-0515

Nguyen MM, Stone MC, Rolls MM. 2011. Microtubules are organized independently of the centrosome in Drosophila neurons. Neural Dev 6:38. doi:10.1186/1749-8104-6-38

Nirschl JJ, Ghiretti AE, Holzbaur ELF. 2017. The impact of cytoskeletal organization on the local regulation of neuronal transport. Nat Rev Neurosci 18:585–597. doi:10.1038/nrn.2017.100

Norkett R, Castillo UD, Lu W, Gelfand VI. 2020. Ser/Thr kinase Trc controls neurite outgrowth in Drosophila by modulating microtubule-microtubule sliding. Elife 9:e52009. doi:10.7554/elife.52009

Ori-McKenney KM, Jan LY, Jan Y-N. 2012. Golgi outposts shape dendrite morphology by functioning as sites of acentrosomal microtubule nucleation in neurons. Neuron 76:921 930. doi:10.1016/j.neuron.2012.10.008

Paz J, Lüders J. 2018. Microtubule-Organizing Centers: Towards a Minimal Parts List. Trends Cell Biol 28:176–187. doi:10.1016/j.tcb.2017.10.005

Petry S, Groen AC, Ishihara K, Mitchison TJ, Vale RD. 2013. Branching Microtubule Nucleation in Xenopus Egg Extracts Mediated by Augmin and TPX2. Cell 152:768–777. doi:10.1016/j.cell.2012.12.044

Popov AV, Severin F, Karsenti E. 2002. XMAP215 Is Required for the Microtubule-Nucleating Activity of Centrosomes. Curr Biol 12:1326–1330. doi:10.1016/s0960-9822(02)01033-3

Qu X, Kumar A, Blockus H, Waites C, Bartolini F. 2019. Activity-Dependent Nucleation of Dynamic Microtubules at Presynaptic Boutons Controls Neurotransmission. Curr Biol 29:4231–4240.e5. doi:10.1016/j.cub.2019.10.049

Rao AN, Patil A, Black MM, Craig EM, Myers KA, Yeung HT, Baas PW. 2017. Cytoplasmic Dynein Transports Axonal Microtubules in a Polarity-Sorting Manner. Cell Reports 19:2210–2219. doi:10.1016/j.celrep.2017.05.064

Rolls MM, Jegla TJ. 2015. Neuronal polarity: an evolutionary perspective. J Exp Biology 218:572–580. doi:10.1242/jeb.112359

Roostalu J, Cade NI, Surrey T. 2015. Complementary activities of TPX2 and chTOG constitute an efficient importin-regulated microtubule?nucleation module. Nat Cell Biol 17:1422 1434. doi:10.1038/ncb3241

Sampaio P, Rebollo E, Varmark H, Sunkel CE, González C. 2001. Organized microtubule arrays in γ-tubulin-depleted Drosophila spermatocytes. Current biology: CB 11:1788 1793. doi:10.1016/s0960-9822(01)00561-9

Sanchez AD, Feldman JL. 2016. Microtubule-organizing centers: from the centrosome to non-centrosomal sites. Curr Opin Cell Biol 44:93 101. doi:10.1016/j.ceb.2016.09.003

Sánchez-Huertas C, Freixo F, Viais R, Lacasa C, Soriano E, Lüders J. 2016. Non-centrosomal nucleation mediated by augmin organizes microtubules in post-mitotic neurons and controls axonal microtubule polarity. Nat Commun 7:12187. doi:10.1038/ncomms12187

Satoh D, Suyama R, Kimura K, Uemura T. 2012. High-resolution in vivo imaging of regenerating dendrites of Drosophila sensory neurons during metamorphosis: local filopodial degeneration and heterotypic dendrite-dendrite contacts. Genes Cells 17:939–951. doi:10.1111/gtc.12008

Scholey JM. 2012. Kinesin-2: A Family of Heterotrimeric and Homodimeric Motors with Diverse Intracellular Transport Functions. Annu Rev Cell Dev Bi 29:443–469. doi:10.1146/annurev-cellbio-101512-122335

Sears JC, Broihier HT. 2016. FoxO regulates microtubule dynamics and polarity to promote dendrite branching in Drosophila sensory neurons. Dev Biol 418:40 54. doi:10.1016/j.ydbio.2016.08.018

Shimono K, Fujimoto A, Tsuyama T, Yamamoto-Kochi M, Sato M, Hattori Y, Sugimura K, Usui T, Kimura K, Uemura T. 2009. Multidendritic sensory neurons in the adult Drosophila abdomen: origins, dendritic morphology, and segment- and age-dependent programmed cell death. Neural Dev 4:37. doi:10.1186/1749-8104-4-37

Slep KC. 2009. The role of TOG domains in microtubule plus end dynamics. Biochem Soc T 37:1002–1006. doi:10.1042/bst0371002

Slep KC, Vale RD. 2007. Structural Basis of Microtubule Plus End Tracking by XMAP215, CLIP-170, and EB1. Mol Cell 27:976–991. doi:10.1016/j.molcel.2007.07.023

Stiess M, Bradke F. 2011. Neuronal polarization: the cytoskeleton leads the way. Dev Neurobiol 71:430 444. doi:10.1002/dneu.20849

Stiess M, Maghelli N, Kapitein LC, Gomis-Rüth S, Wilsch-Bräuninger M, Hoogenraad CC, Tolić-Nørrelykke IM, Bradke F. 2010. Axon Extension Occurs Independently of Centrosomal Microtubule Nucleation. Science 327:704–707. doi:10.1126/science.1182179

Tanaka N, Meng W, Nagae S, Takeichi M. 2012. Nezha/CAMSAP3 and CAMSAP2 cooperate in epithelial-specific organization of noncentrosomal microtubules. Proc National Acad Sci 109:20029 20034. doi:10.1073/pnas.1218017109

Tas RP, Chazeau A, Cloin BMC, Lambers MLA, Hoogenraad CC, Kapitein LC. 2017. Differentiation between Oppositely Oriented Microtubules Controls Polarized Neuronal Transport. Neuron 96:1264–1271.e5. doi:10.1016/j.neuron.2017.11.018

Thawani A, Kadzik RS, Petry S. 2018. XMAP215 is a microtubule nucleation factor that functions synergistically with the γ-tubulin ring complex. Nat Cell Biol 20:1 18. doi:10.1038/s41556-018-0091-6

Tillery M, Blake-Hedges C, Zheng Y, Buchwalter R, Megraw T. 2018. Centrosomal and Non-Centrosomal Microtubule-Organizing Centers (MTOCs) in Drosophila melanogaster. Cells 7:121. doi:10.3390/cells7090121

Tovey CA, Conduit PT. 2018. Microtubule nucleation by γ-tubulin complexes and beyond. Essays Biochem 91:EBC20180028. doi:10.1042/ebc20180028

Tovey CA, Tubman CE, Hamrud E, Zhu Z, Dyas AE, Butterfield AN, Fyfe A, Johnson E, Conduit PT. 2018. γ-TuRC Heterogeneity Revealed by Analysis of Mozart1. Curr Biol 28:2314 2323.e6. doi:10.1016/j.cub.2018.05.044

Tsuchiya K, Goshima G. 2021. Microtubule-associated proteins promote microtubule generation in the absence of γ-tubulin in human colon cancer cells. Biorxiv 2021.08.13.456214. doi:10.1101/2021.08.13.456214

Valenzuela A, Meservey L, Nguyen H, Fu M. 2020. Golgi Outposts Nucleate Microtubules in Cells with Specialized Shapes. Trends Cell Biol. doi:10.1016/j.tcb.2020.07.004

Vinogradova T, Miller PM, Kaverina I. 2009. Microtubule network asymmetry in motile cells: role of Golgi-derived array. Cell Cycle 8:2168 2174. doi:10.4161/cc.8.14.9074

Wang S, Wu D, Quintin S, Green RA, Cheerambathur DK, Ochoa SD, Desai A, Oegema K. 2015. NOCA-1 functions with γ-tubulin and in parallel to Patronin to assemble non-centrosomal microtubule arrays in C. elegans. Elife 4:e08649. doi:10.7554/elife.08649

Wang Y, Rui M, Tang Q, Bu S, Yu F. 2019. Patronin governs minus-end-out orientation of dendritic microtubules to promote dendrite pruning in Drosophila. Elife 8:e39964. doi:10.7554/elife.39964

Weiner AT, Lanz MC, Goetschius DJ, Hancock WO, Rolls MM. 2016. Kinesin-2 and Apc function at dendrite branch points to resolve microtubule collisions. Cytoskeleton 73:35–44. doi:10.1002/cm.21270

Weiner AT, Seebold DY, Torres-Gutierrez P, Folker C, Swope RD, Kothe GO, Stoltz JG, Zalenski MK, Kozlowski C, Barbera DJ, Patel MA, Thyagarajan P, Shorey M, Nye DMR, Keegan M, Behari K, Song S, Axelrod JD, Rolls MM. 2020. Endosomal Wnt signaling proteins control microtubule nucleation in dendrites. Plos Biol 18:e3000647. doi:10.1371/journal.pbio.3000647

Wieczorek M, Bechstedt S, Chaaban S, Brouhard GJ. 2015. Microtubule-associated proteins control the kinetics of microtubule nucleation. Nat Cell Biol 17:907 916. doi:10.1038/ncb3188

Williams DW, Truman JW. 2005. Cellular mechanisms of dendrite pruning in Drosophila: insights from in vivo time-lapse of remodeling dendritic arborizing sensory neurons. Development 132:3631–3642. doi:10.1242/dev.01928

Woodruff JB, Gomes BF, Widlund PO, Mahamid J, Honigmann A, Hyman AA. 2017. The Centrosome Is a Selective Condensate that Nucleates Microtubules by Concentrating Tubulin. Cell 169:1066 1077.e10. doi:10.1016/j.cell.2017.05.028

Wu J, de Heus C, Liu Q, Bouchet BP, Noordstra I, Jiang K, Hua S, Martin M, Yang C, Grigoriev I, Katrukha EA, Altelaar AFM, Hoogenraad CC, Qi RZ, Klumperman J, Akhmanova A. 2016. Molecular Pathway of Microtubule Organization at the Golgi Apparatus. Dev Cell 39:44 60. doi:10.1016/j.devcel.2016.08.009

Yalgin C, Ebrahimi S, Delandre C, Yoong LF, Akimoto S, Tran H, Amikura R, Spokony R, Torben-Nielsen B, White KP, Moore AW. 2015. Centrosomin represses dendrite branching by orienting microtubule nucleation. Nat Neurosci 18:1437 1445. doi:10.1038/nn.4099

Yan J, Chao DL, Toba S, Koyasako K, Yasunaga T, Hirotsune S, Shen K. 2013. Kinesin-1 regulates dendrite microtubule polarity in Caenorhabditis elegans. Elife 2:e00133. doi:10.7554/elife.00133

Yang SZ, Wildonger J. 2020. Golgi Outposts Locally Regulate Microtubule Orientation in Neurons but Are Not Required for the Overall Polarity of the Dendritic Cytoskeleton. Genetics 215:435–447. doi:10.1534/genetics.119.302979

Yau KW, Schätzle P, Tortosa E, Pagès S, Holtmaat A, Kapitein LC, Hoogenraad CC. 2016. Dendrites In Vitro and In Vivo Contain Microtubules of Opposite Polarity and Axon Formation Correlates with Uniform Plus-End-Out Microtubule Orientation. J Neurosci 36:1071–1085. doi:10.1523/jneurosci.2430-15.2016

Zheng Y, Buchwalter RA, Zheng C, Wight EM, Chen JV, Megraw TL. 2020. A perinuclear microtubule-organizing centre controls nuclear positioning and basement membrane secretion. Nat Cell Biol 22:297–309. doi:10.1038/s41556-020-0470-7

Zheng Y, Buchwalter RA, Zheng C, Wight EM, Chen JV, Megraw TL. 2019. A perinuclear microtubule-organizing center controls nuclear positioning and basement membrane secretion. Biorxiv 2019.12.24.888065. doi:10.1101/2019.12.24.888065

Zhou W, Chang J, Wang X, Savelieff MG, Zhao Y, Ke S, Ye B. 2014. GM130 is required for compartmental organization of dendritic golgi outposts. Curr Biol 24:1227 1233. doi:10.1016/j.cub.2014.04.008

Zupa E, Liu P, Würtz M, Schiebel E, Pfeffer S. 2021. The structure of the γ-TuRC: a 25-years-old molecular puzzle. Curr Opin Struc Biol 66:15–21. doi:10.1016/j.sbi.2020.08.008

